# The Molecular Picture of the Local Environment in a Stable Model Coacervate

**DOI:** 10.1101/2024.01.12.575416

**Authors:** Atanu Baksi, Hasan Zerze, Aman Agrawal, Alamgir Karim, Gül H. Zerze

**Affiliations:** William A. Brookshire Department of Chemical and Biomolecular Engineering, University of Houston, Houston, Texas 77204, United States; Department of Chemistry and Pritzker School of Molecular Engineering, University of Chicago, Chicago, IL 60637

**Keywords:** COACERVATE STABILITY, LOCAL IONIC ENVIRONMENT, POLYELECTROLYTES, ION RESIDENCE TIMESCALE

## Abstract

Polymers with electric charge, known as polyelectrolytes, are well known to form complex coacervates, which have vital implications in various biological processes and beyond. While significant advancements have been made in comprehending the molecular interactions that *drive* complex coacervation, the interactions that *stabilize* the coacervates against coalescence present an intricate experimental challenge and remain a subject of ongoing investigation. In a recent experimental study, polydiallyldimethylammonium chloride polycationic (PDDA) and anionic adenosine triphosphate (ATP) coacervates have been shown to stabilize upon transferring them to deionized water. Here, we perform molecular dynamics simulations of PDDA-ATP coacervates both in supernatant and in DI water, to understand the ion dynamics and structure within stable coacervates. We produced and analyzed an aggregated sum of 63 *μs* simulation data of PDDA-ATP coacervates in explicit water when they are in supernatant and deionized (DI) water. We found that discarding the supernatant and transferring the coacervates to DI water causes an immediate ejection of a significant amount (more than 50%) of small ions (*Na*^+^ and *Cl*^−^) from the coacervates to the bulk solution. Subsequently, the DI water environment alters the ionic density profiles in coacervates and the surface ion dynamics. We calculated a notable slowdown for the coacervate ions when they were transferred to the DI water. These results suggest that the initial ejection of the ions from the coacervates in DI water potentially brings the outer layer of the coacervates to a physically bound state that prevents or slows down the further mobility of ions.

**Significance Statement:** Complex coacervates are promising agents for encapsulating and delivering various materials in living organisms, however, they are often prone to coalesce, limiting the range of their applications. Recently, these coacervates have been stabilized by transferring them to deionized water. However, a molecular understanding of this stability against coalescence remained elusive. This study utilizes computer simulations to model a stable coacervate system previously probed experimentally. When the coacervates were transferred to deionized water, a significant portion of the ions were immediately ejected into the solution, modifying the coacervates’ total charge and facilitating formation of possible surface crust. These molecular insights into the stable coacervates will enable their controllable design for encapsulation and delivery applications.

---

Solutions of two oppositely charged macromolecules have been observed to separate into two liquid-like phases with different viscosities, commonly referred to as coacervation/condensation as originally described by Bunderberg de Jong and Kruyt for colloidal mixtures (1) while the history of biomolecular condensates dates back nearly a century earlier (2–4). The polyelectrolyte complex coacervates (PECCs) play a vital role in biological contexts e.g., intracellular organization (5–12) and early stages of prebiotic evolution (6, 13, 14). Moreover, numerous applications of PECCs including their usage in food science applications (15, 16) biomimetic adhesives (17), and advanced underwater adhesives (18, 19) spurred a growing interest in polyelectrolyte coacervates. Peptide-based coacervates have novel therapeutic applications, such as tissue regeneration and drug delivery systems (20). Furthermore, efforts have been made to design controllable biomimetic field-responsive soft particles in the form of coacervates, which might serve as vehicles for transporting various reactive species related to synthetic drugs or genetic materials (21).

As these versatile coacervates are adapted to lead more and more promising applications, a fundamental understanding of the interactions stabilizing the coacervates and driving the formation of these coacervates has become crucial to understand. Accordingly, in addition to the efforts in advancing applications of PECCs, an appreciable amount of effort has been dedicated to studying their fundamental biophysical properties experimentally (22–33), shedding light on fundamentals of coacervates including their thermodynamic stability, phase coexistence behavior, rheological properties, molecular partitioning. Moreover, theoretical (34–42) and computational (43–48) works focused on determining molecular determinants of complex coacervation and phase behavior of coacervates, predicting interfacial tension of coacervates, and explaining different thermodynamic contributors in coacervation process. More recent efforts in the literature have focused on producing thermally and chemically stable coacervates (21, 49–51). In earlier work, Williams et al. (49) have reported that high ionic strength discourages the PDDA-ATP coacervation and emphasized the significance of charge interaction in droplet formation and stability. They have further found that the formed droplets spontaneously become positively or neutrally charged depending on the molar PDDA monomer to ATP ratio. Later, Karim and coworkers have experimentally produced coacervates that do not coalesce over time into a macrophase (21). They have produced these stable PECCs by transferring them to a deionized environment, which presumably stabilizes their otherwise highly diffuse interfaces. Despite this significant shift in the stability of PECCs (without requiring additives or chemical reactions), the underlying molecular picture leading to this stability has remained not well understood. Although the above-mentioned computational and theoretical studies have been instrumental in delineating the phase equilibrium and rheological properties of coacervates, the large (typically, prohibitively large) costs of explicit solvation and related system size requirements limited the molecular dynamics (MD) simulation studies to modeling polyelectrolytes in implicit solvents (52–57). Consequently, a molecular picture of changes in the local solvation environment (including explicit water and ions), dynamics, and structure stabilizing the complex coacervates have remained crucially needed, as it would allow the development of novel tools to control and tune the stability and properties of PECCs.

In this study, we intended to gain an understanding of the effect of local ionic solution environments in stable coacervates at a molecular level with the explicit presence of water and ions. Accordingly, we employed explicit-solvent coarse-grained (CG) MD simulations to study coacervates of polycationic polydiallyldimethylammonium chloride (PDDA) polymers and anionic adenosine triphosphate (ATP) molecules in explicit water and counterions. Subsequently, the identity of the ionic supernatant environment was altered to deionized (DI) water (which is analogous to transferring the coacervates into DI water as in the experimental study (21)) to investigate the impact of this deionized environment on the coacervate structure and ion dynamics. We found that the PDDA-ATP coacervates are positively charged and they become more positively charged after transferring them to DI water. Importantly, we also found that a significant fraction of the small ions (*Na*^+^ and *Cl*^−^) within the coacervates are immediately ejected into the ion-deprived bulk solution when the coacervates are transferred to DI water with a subsequent slowdown in the ion dynamics for the ions remaining in the coacervates. The majority of the ejected ions used to reside on the surface of the coacervates before transferring them to the DI water. These findings led us to conclude that the initial ejection of the ions (most of which are ejected from the surface) contributes to a hypothetical crust-like surface formation in DI water, which also explains our measurement of the subsequent slowdown of ions. While the crust-like surface layer formation in DI water has originally been hypothesized by Karim and coworkers (21), this work provides a direct quantification of electrostatic changes upon DI water transfer lending further support to the crust-like layer formation hypothesis and marks the first extensive investigation of molecular picture of the electrostatic changes in stable coacervates.

## Methods

We have employed MARTINI2.0 force field parameters for simulating PDDA-ATP coarcervates. To ensure adequate statistical significance and reproducibility of our findings we have performed 8 independent simulations starting from randomly monodispersed mixtures of PDDA and ATP. Formation of the coacervate from the dispersed PDDA and ATP molecules in an aqueous medium can be discernable through the time evolution of PDDA and ATP fraction in the cluster in Figure 1A. To analyze the region-specific ion dynamics, we defined the regions in the coacervates projecting PDDA and ATP positions on three principal planes; an example of which is shown in Figure 1B for a selected coacervate. Analogous plots for all 8 independent simulations (*Supporting Information (SI) Appendix, Figures S1 - S6*), details of the modeling and force field, simulation parameter details, and theoretical background for analyses carried out in this study can be found in the *SI Appendix, Text SI. 1(a)-1(e)*.

**Fig. 1.**
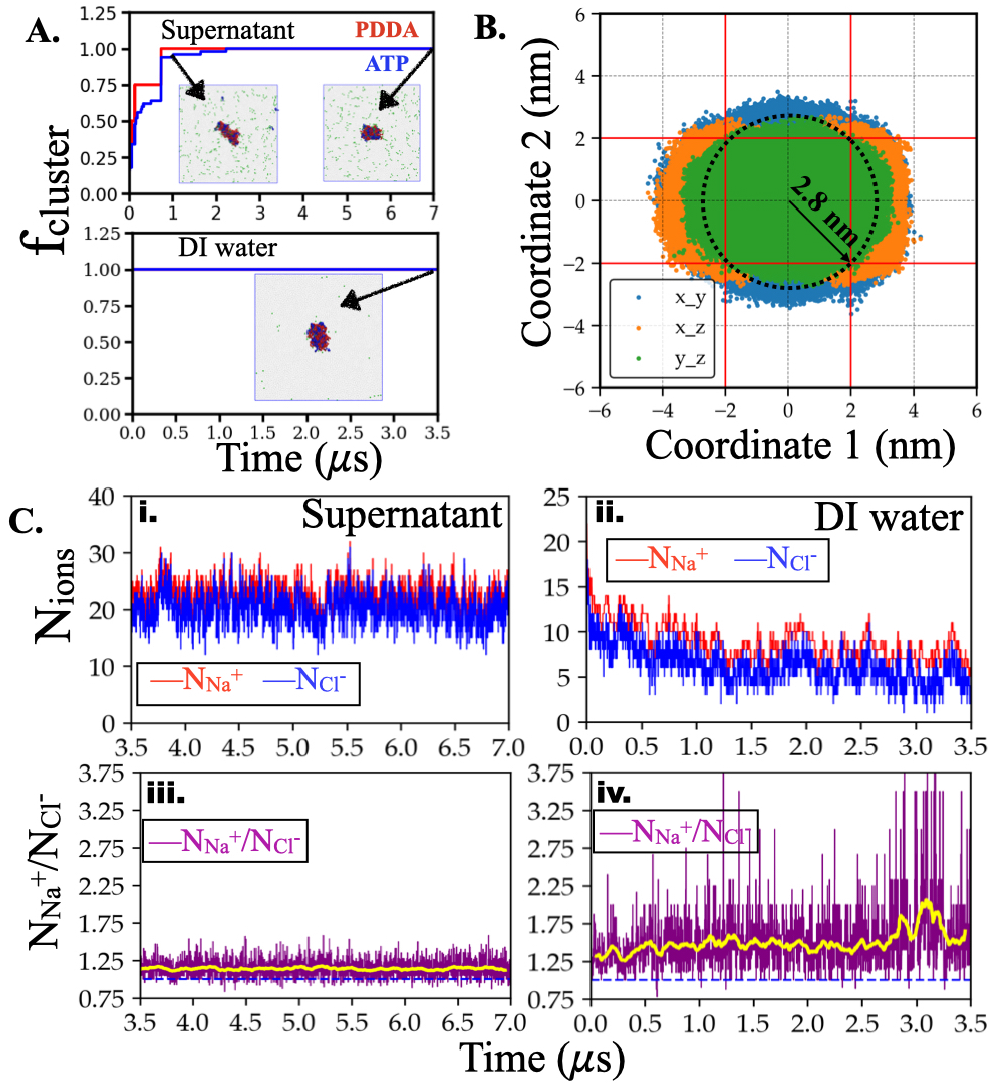
Coacervate formation, coacervate surface and core definitions, and the ionic nature of the coacervates. A. Time evolution of the fraction of PDDA and ATP molecules in the largest cluster (*f*_*cluster*_) in ‘supernatant’ (top) and in ‘DI water’ (bottom). We consider the cluster to be fully formed after *f*_*cluster*_ reaches a stable value of 1.0. The snapshots in the insets are the simulation box at indicated simulation times where PDDA, ATP, and small ions beads are colored red, blue, and green, respectively. B. Projection of the clustered PDDA and ATP beads’ (coacervates) positions on three principle planes of the cluster. This projection informs about the size and the shape (e.g., asphericity) of the coacervates. We also used these projections to label the ‘surface’ and ‘core’ of coacervates for the selected simulation. The overlapping rectangular area is used to identify a core represented by a sphere with a radius of *r*_*core*_=2.8 nm. In between *r*_*core*_=2.8 nm and *r*_*surface*_=5 nm is considered as surface. Here, coordinate 1 and 2 are the pairs of principal axes orthogonal to the projected axis. Three colored regions are projections of PDDA and ATP beads on three principal planes. C. Time evolution of the number of *Na*^+^ and *Cl*^−^ ions within the coacervate when it is in ‘supernatant’ (i) and ‘DI water’ (ii) for a selected set. Time evolution of the ratio of the number of *Na*^+^ to *Cl*^−^ ions within the coacervate in ‘supernatant’ (iii) and ‘DI water’ (iv). Yellow lines in the lower panel indicate a running average of the ratio of the number of *Na*^+^ to *Cl*^−^ ions, which is higher than 1 both in supernatant and DI water.

## Results and Discussions

### Ionic nature of the cluster

We first quantified the ion distribution in coacervates both in ionic supernatant and in DI water (i.e., after transferring the coacervates to DI water). For that, we counted the number of *Na*^+^ and *Cl*^−^ ions within the coacervates. Since all of our PDDA and ATP molecules are clustered in the coacervates (i.e., no PDDA or ATP in the bulk solution), the ions that are at a distance smaller than 1 nm to the PDDA and ATP beads are considered to be in the coacervates. Figure 1C depicts the time evolution of ion numbers within the cluster (upper panel) and the ratio of the positive (*Na*^+^) to negative (*Cl*^−^) ions (lower panel) both for the cases of supernatant (i, iii) and DI water (ii, iv) for a selected simulation. The ratio is consistently observed to be greater than 1.0 in both ‘supernatant’ and ‘DI water’ for all the independent simulations (Figure S7 and Figure S8, respectively). Given that PDDA and ATP molecules have equal and opposite charges and are always clustered within the coacervates (i.e., no PDDA or ATP in bulk solution), the *Na*^+^/*Cl*^+^ ratio being larger than 1 means the coacervates are positively charged.

We have observed that the number of *Na*^+^ and *Cl*^−^ ions within the coacervates initially decreases as a function of simulation time after the coacervate was transferred to DI water as seen in Figure 1C(ii) while *Na*^+^ to *Cl*^−^ ratio increases (Figure 1C lower panels’ comparison, and Figure S7 and Figure S8 panels’ comparison). The former is a result of the concentration gradient of ions that favors the movement of ions towards the initially ion-free bulk phase. We note that the number of *Na*^+^ and *Cl*^−^ ions maintain equilibrium after 1.5*μs*. We found that the *Na*^+^ to *Cl*^−^ number ratio, slightly increases almost immediately, after placing the coacervate into DI water (see Figure 1C(iv)). Since all PDDA and ATP molecules are clustered in the coacervates all the time, the increased Na+ predominance in the counterions inside the coacervates indicates that coacervates become even more positively charged when transferred in DI water.

### Density profile and size of the cluster

We also analyzed the density of the PDDA, ATP, counterions, and water molecules within the coacervates (Figure 2A). PDDA and ATP density profiles showed that the PDDA-ATP clusters both in the supernatant and DI water extend up to a radius of about 5 nm from the center of the cluster (Figure 2A i. and ii.). As a guide to the eye, we indicated this 5 nm radius with vertical dashed lines in Figure 2A but we note that this vertical line does not necessarily define the surface boundary of the coacervates due to the deviations from a spherical shape at this length scale as discussed in the next section.

**Fig. 2.**
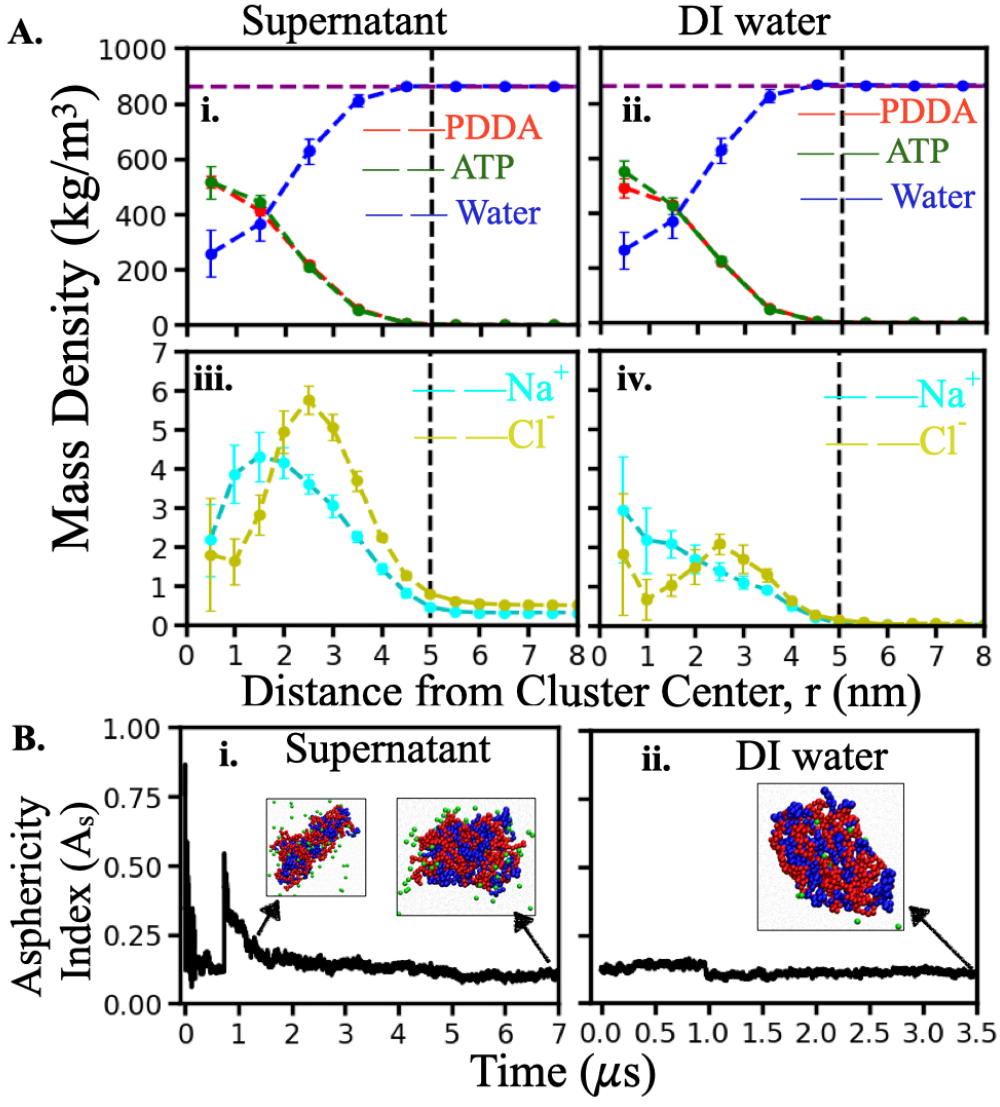
Density profiles and asphericity of the coacervates. A. Mass density profiles of PDDA, ATP, water, and ions in supernatant (left panels) and DI water (right panels). Error bars are the standard deviation of sample means. The density profiles are averaged over eight independent simulations. The vertical line at 5 nm is the guiding line indicating the maximum extent of the cluster. The horizontal line at 862 kg/m^3^ refers to equilibrated MARTINI 2.0 water density at 298 K. The molecular weight of each PDDA monomer in its fully dissociated state is 126.21 g/mol and that of an ATP molecule at the protonation state of −4*e* is 503.2 g/mol. The molecular weight of water, *Na*^+^ and *Cl*^−^ ions are 18.01, 22.99, and 35.45 g/mol, respectively. B.Time evolution of asphericity index of the largest cluster in (i) supernatant (ii) DI water. Attached snapshots in the inset are the clusters at indicated simulation times where PDDA, ATP, and ion beads are colored in red, blue, and green, respectively.

We found that the coacervates contain a significant amount of water and ions after transferring them to DI water in agreement with their liquid-like nature. We note that the water density plateaus at around 862 kg/m^3^, which is consistent with the bulk density of this water model at these thermodynamic conditions (Figure S9). Since our cluster size is relatively small, PDDA and ATP densities inside the coacervate do not have a clear plateau, however, the data were sufficient to produce sigmoidal fits (Figure S10). From the sigmoidal fits, we predict the average PDDA and ATP densities in the coacervates (*C*_*dense*_) as approximately 540 kg/*m*^3^ and 536 kg/*m*^3^, respectively, in the supernatant. In DI water, *C*_*dense*_ for PDDA is approximately 511 kg/*m*^3^ and that for ATP is approximately 566 kg/*m*^3^ (Figure S10).

In addition, we estimated the average water content (mass fraction) within the coacervates by calculating the total mass of water molecules present within the coacervate (water molecules that are at a distance smaller than 1 nm to the PDDA and ATP beads) and normalizing it with the total mass of the clustered molecules (PDDA, ATP, ions, and water). We found that the average water content within the coacervates is approximately 60% both in the supernatant and DI water systems, which is close to the experimental measurements of 70% and 65% (by mass) of water in the supernatant and DI water, respectively (21).

We found that the mass density profiles of PDDA and ATP within the coacervates are similar to each other, for both supernatant and DI water systems, with slightly larger ATP concentrations at the center for both systems (Figure 2A, top panels). Note that this slight difference in concentration is amplified by the four-fold difference in ATP (−4e) and PDDA monomer (+1e) valence, making the core of the coacervates more negatively charged. We also visually detected that ATP occupies a larger portion of the center (Movie S1), which becomes more pronounced after transferring the coacervates to DI water. We also found that *Cl*^−^ ions mostly occupy the surface of the coacervates whereas the *Na*^+^ ions populate more evenly at the center of the supernatant coacervates (Figure 2A, bottom panels). The slightly larger concentration of ATP at the coacervate center is consistent with the larger *Na*^+^ concentration at the center.

While the macroion density profiles remain similar, two important changes happen to the counterion density profiles upon transferring the coacervates to DI water. i) Ion densities in the coacervates drastically drop after they are placed in DI water. The majority of the ions from the coacervates are ejected into the bulk solution after being placed in DI water (Figure 1C. ii., also see *SI Appendix, Movie S1*). ii) *Na*^+^ ion density peaks more sharply at the center after DI water placement (Figure 2A iii. and iv.). In both supernatant and DI water, the core region of the coacervates is populated more by *Na*^+^ ions and the difference between *Na*^+^ and *Cl*^−^ density profiles becomes more pronounced after DI water placement. *Na*^+^ ion density profile sharply peaks at the center with a monotonic decrease as the distance away from the center increases in DI water while the *Cl*^−^ density trend remains similar in DI water to that in supernatant. The *Na*^+^ ions being buried at the center of the coacervates significantly affects the ion dynamics within the coacervates, which we quantified in *Ions dynamics within coacervates* subsection.

Since the density profiles provide limited information about the shape of coacervates, we calculated the asphericity index, *A*_*s*_, of the coacervates. Instantaneous *A*_*s*_ of coacervates is calculated following *SI Appendix, eqn S1*) as a function of time. Figure 2B shows the time evolution of *A*_*s*_ of the largest cluster for both (i) supernatant and (ii) DI water systems for one of the simulation sets with the snapshots of the coacervates collected at indicated simulation times. *A*_*s*_ and the representative snapshots of the coacervates for all independent simulations are presented in *SI Appendix, Figure S11 and S12* for supernatant and DI water systems, respectively. These results show that the asphericity index of the well-stable coacervates (after simulation time 3.5 μs) varies within 0.1 *< A*_*s*_ *<* 0.4 for different independent simulations. We observe after transferring the clusters to DI water, coacervate morphology exhibits minimal variation, characterized by a low degree of fluctuation. This observation is consistent with the higher stability of the coacervates against dissociation in DI water as compared to ‘supernatant’ systems.

### Ion dynamics within coacervates

To further extend our understanding of the coacervate stability in DI water, we analyzed the dynamics of the counterions within the coacervate surface and core, both in supernatant and in DI water. As our coacervates are not perfectly spherical (see the subsection above), we first identified the ‘core’ and ‘surface’ regions in our coacervates based on the principal axes projection (Figure 1B). For the full protocol of the principle axes projection and labeling, see the *SI Appendix, Text SI. 1(e)*.

#### Time evolution of the location of tagged ions after transferring the coacervates to DI water

Following the definition of the ‘core’ and ‘surface’ regions, we tagged the *Na*^+^ and *Cl*^−^ ions as initially-core and initially-surface ions at the time of transferring the coacervates to DI water. We then followed the displacement of tagged ions from their initially tagged region (surface or core) to other (surface, core, or bulk solvent) regions and calculated the fraction of initially tagged ions in those regions throughout the simulation time. Figure 3A shows the fluctuations as a function of time for one of the simulations, where the upper panel depicts the time evolution of the fraction of initially surface ions that stays on the surface (green) or moves to bulk solvent (red) or core (black). Figure 3B represents the time evolution of the fraction of initially-core ions that stay in the core (black), and move to the surface (green) or to bulk solvent (red). We find the ions from the coacervates are rapidly ejected in the ion-deprived DI water solvent as can be seen in the bottom panels of Figure 3A and 3B. A schematic diagram of the rapid ejection of these initially tagged ions is presented in Figure 3C. Over time, we calculated that ∼ 75% of the initially-surface ions move to the bulk region outside the cluster, and the remaining (∼ 25%) ions remain inside the cluster (core or surface region). On the other hand, only ∼ 60% of the initially-core ions are found to move to the bulk region and the remaining initially-core ions stay within the coacervate (in the core or the surface).

**Fig. 3.**
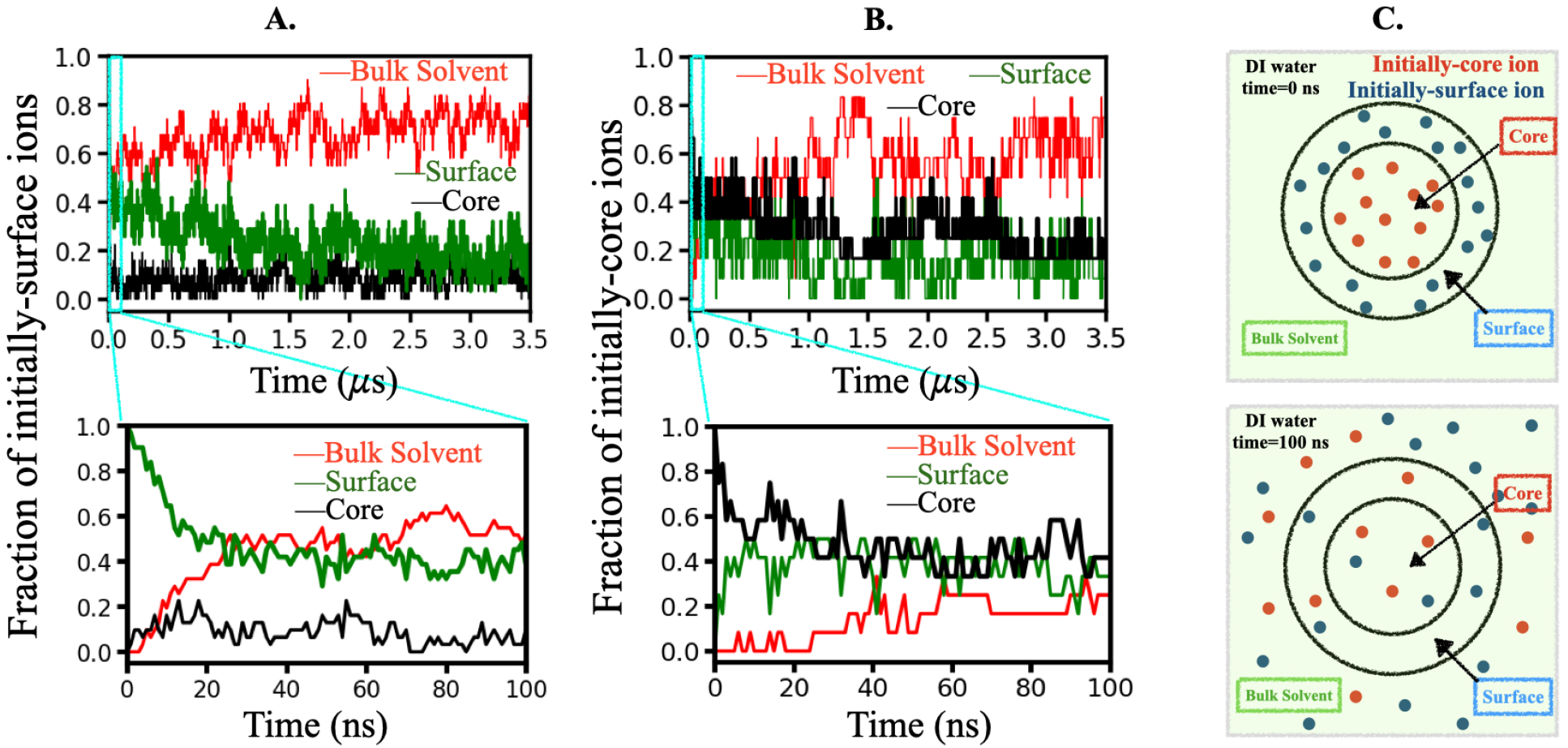
Signature of rapid ejection of ions from coacervates upon transferring them to DI water. The ions located within a spherical region of radius *r*_*core*_=2.8 nm are labeled as core ions whereas the ions located in the spherical region between *r*_*core*_=2.8 nm and *r*_*surface*_=5 nm labeled as surface ions at t=0 ns. Results are shown for one simulation set. Results for other sets are shown in the SI Appendix, Figure S6). Ion ejection from the surface (A.) and from the core (B.). The top panels in A and B show the time evolution of the label change of the ions initially labeled as ‘surface’ and the time evolution of the label change of the ions initially labeled as ‘core’, respectively, after coacervates transferred to DI water. The initial 100 ns of the simulation data is zoomed in on the bottom panels of A and B to show the initial rapid ejection both from the surface (A.) and from the core (B.). C. Schematic diagram of the ejection of initially-core and initially-surface ions.

These findings indicate that a large percentage of both the initially-surface and initially-core ions transition to the bulk solution but nearly 25% more of the initially-surface ions will transition to bulk solvent compared to initially-core ions. We observed a similar trend in all eight simulations in DI water (shown in *SI Appendix, Figures S13 and S14*). We also calculated the total fraction of ions ejected into the DI water. We found that on average ∼ 65% of the ions ejected into DI water come from the coacervate surface whereas ∼ 35% come from the coacervate core. This observation also supports the possibility of a crust-like layer formation at the coacervate surface in DI water, blocking the mobility of ions in the coacervate core.

#### Ion residence times depend on the location in the coacervate

We then calculated the ion residence times, from a time correlation function, 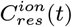, for the ions residing in the core and the surface regions, for both in the supernatant and DI water surroundings following eqn S2, and eqn S3 (see SI Appendix Text SI. 1e for the details of the calculation). The correlation function decays are presented in *SI Appendix, Figures S15 - S20*. Furthermore, to qualitatively understand contributions from oppositely charged counterions on total ion dynamics, we also separately analyze the correlation decay of *Na*^+^ and *Cl*^−^ ions, individually. Figures S15 (for supernatant) and S16 (for DI water) show the time-dependent decay of the residence time correlation function for all ions collectively (*Na*^+^ and *Cl*^−^ ions combined); Figures S17 (for supernatant) and S18 (for DI water) show the same for positively charged *Na*^+^ ions only; and Figures S19 (supernatant) and S20 (DI water) show the same for negatively charged *Cl*^−^ ions only.

To be able to calculate the timescales associated with these correlation decays, namely residence timescales, we fit these time correlation functions using multi-exponential fit-functions as follows:

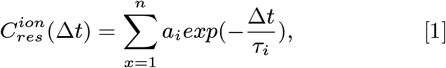

where *τ*_*i*_ are the relaxation (residence) time scales, *a*_*i*_ are prefactors, and the index *i* runs over 2 or 3 depending on whether the bi-exponential or tri-exponential function represents data better. Fits are presented together with raw data on the plots from *SI Appendix, Figures S15 - S20*.

We found that the correlation decays for all the ions residing in the ‘core’ regions from all the independent simulations can be adequately described with three distinctive characteristic timescales, 1) of the order of nanoseconds, *τ*_*fast*_ *<* 10 *ns*, 2) of the order of tens of nanoseconds (10 *ns < τ*_*intermediate*_ *<* 200 *ns*), and 3) of the order of hundreds of nanoseconds to tens of microseconds (200 *ns < τ*_*long*_ *<* 50 *μs*). On the other hand, for all ions residing in the ‘surface’ region, the correlation decays exhibit mainly two characteristic timescales, which were previously identified as the first two timescales describing ‘core’ ion dynamics, *τ*_*fast*_ and *τ*_*intermediate*_. The fast timescale (*τ*_*fast*_) can be attributed to the rapid exchange of ions between different regions, whereas the intermediate timescale (*τ*_*intermediate*_) is associated with the translational motion of ions within those specific regions. The slowest characteristic timescales *τ*_*long*_ are indicative of kinetically trapped ions, compelling them to remain in the particular region for an extended duration.

Characteristic timescales observed from different independent simulations are summarized in *SI Appendix, Tables S1 S12*. The timescales (with associated amplitude) that are averaged over all independent simulations and their standard deviations (as errors) are reported in Figure 4 in log scale. The left panels in Figure 4 represent residence timescales of ions residing in the ‘core’ region whereas the right panels depict the same for ions residing in the ‘surface’ region. The top panels show the characteristic timescales describing correlation decays with contribution from all ions, whereas the middle and bottom panels show the same for only *Na*^+^ ions and only *Cl*^−^ ions, respectively. The comparisons between the residence times in supernatant and DI water systems are provided in each panel.

**Fig. 4.**
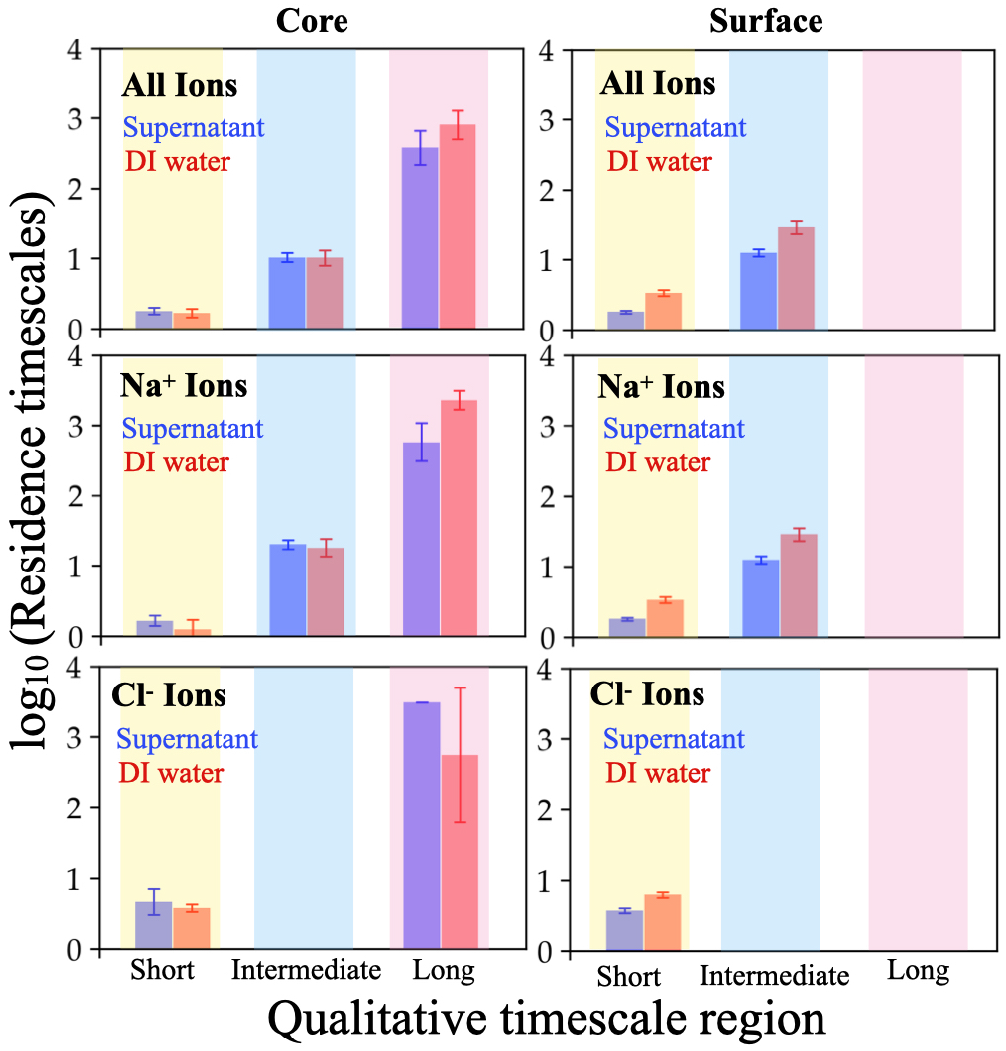
Comparison of characteristic timescales (*log*_10_ (*τ*)) describing ion residence correlation decays in ‘core’ (left panel) and ‘surface’ region (right panel). The top panels report the timescales associated with overall ion dynamics whereas the middle and bottom panels report the same separately for *Na*^+^ ions and *Cl*^−^ ions, respectively. Each plot compares the dynamics of ions in the supernatant and DI water system.

The comparisons from Figure 4 showed three interesting findings:

1. The positively charged *Na*^+^ ions are rate-limiting for ion dynamics, which is consistent with them being buried in the coacervate core. Negatively charged *Cl*^−^ ions are much shorter lived within the coacervate and move relatively faster compared to their positive counterpart, *Na*^+^ ions.
2. Ions residing in the ‘surface’ region are faster compared to ions that are residing in the ‘core’ region. This has been observed systematically over all the independent simulations both in supernatant and DI water systems.
3. Ions, particularly the rate-limiting *Na*^+^ ions within the coacervates, exhibit longer lifetimes in DI water systems compared to those in supernatant systems.

### Discussions and Concluding Remarks

Complex coacervates are highly attractive materials as drug delivery agents, protocellular environments, and model membraneless organelles. However, at thermodynamic equilibrium, small coacervate droplets coalesce to form a condensed macro phase limiting the range of their applications. Recent work by Karim and coworkers showed that it is possible to stabilize coacervates against coalescence in a deionized environment, however, the underlying molecular picture of this stability remained elusive and challenging to obtain experimentally.

How does changing the equilibrium environment of coacervate surroundings affect the dynamics of its components? Our findings show interesting ion dynamics when such a non-equilibrium change is introduced in the form of deprivation of counterions from the supernatant. Counterions within the coacervates were rapidly ejected into bulk solution upon DI water transfer. Importantly, the majority of those ejected ions were ejected from the coacervate surface. Subsequently, the ion residence times became at least an order of magnitude slower within coacervates and ionic density profiled underwent restructuring in DI water whereas the density distribution of PDDA, ATP, and water molecules within the coacervates remained mostly unchanged. Since we do not measure a significant change either in the water content or in the polymer density of the coacervates before and after DI water transfer, we do not have an obvious reason for a viscosity change in the coacervates. Since we eliminate the reasons for a viscosity change, we argue that our findings are consistent with crust-like surface layer formation (slowing down the ions inside coacervates), supporting the original hypothesis by Karim and coworkers (21).

We also consistently quantified that PDDA-ATP coacervates consistently carry a net positive charge in the ionic supernatant throughout all our independent simulations. Moreover, we also quantified that the coacervates became more positively charged after transferring them to the DI water system. This suggests that an electrostatic repulsion could also be an additional factor contributing to the coacervate stability in DI water as like charges repel each other.

We argue that the characteristic charge of the coacervates and the slow counterion (i.e., Na+ or Cl-) is determined by the relative size of the macroions. We observed that the smaller macroion (ATP) often arranges itself as small clusters inside the coacervate (*SI Appendix, Movie S1*) and has a slight preference for the center, which becomes more pronounced upon transferring the coacervates to DI water. The slight increase in the ATP concentration in the center is accompanied by a larger concentration of the oppositely charged Na+ counterion at the center, making Na+ ions slower. We also performed the charge-personality swap simulations, that is, simulations of hypothetical PDDAATP systems where we swap the charges of PDDA and ATP molecules (i.e., PDDA beads were made negatively charged and ATP beads were made positively charged, keeping everything else the same, including identities of the counterions, bonded interactions, van der Waals interactions, etc.). We found that the *Cl*^−^ ions predominate the *Na*^+^ ions in the coacervates in that case (*SI Appendix, Figure S21*.), making them negatively charged. Moreover, *Cl*^−^ becomes more buried in the coacervate core and becomes the slower counterion in that case (*SI Appendix, Figures S21 and S22*.). These findings support our argument that the characteristic charge depends on the relative size of the macroions. The counterion that is oppositely charged to the smaller macroion prefers to occupy the core, becomes the slow ion, and predominates the other counterion in number determining the net charge of the coacervate.

This work shows that the coacervates that are stabilized in DI water have distinct changes to their ionic attributes such as the rapid ejection of small counterions, a subtle change in total net charge, and the change in ion dynamics. We believe that these observations apply to systems with macroions of largely different sizes (e.g., PDDA and ATP) as discussed above. This work marks the first investigation of the molecular picture of the local ionic environment of PECCs which are stable against coalescence, enabling further investigation to produce and understand stable PECCs with various macroion sizes and valence. We note that further investigation is still needed to obtain a more complete picture. One limiting factor in this study was the relatively small size of the system (4 PDDA polymers and 50 ATP molecules), which does not allow quantification of local structuring and polymer arrangements on the surface. Since these simulations are performed in the explicit solvent, bigger systems have prohibitively large costs to sample all the relevant parts of the phase space. A future investigation with appropriate advanced sampling techniques will give us more information about the local polymer conformation on the coacervate surface providing us with a more complete picture of the forces stabilizing the coacervates against coalescence.

## Data, Materials, and Software Availability

All data needed to evaluate the conclusions in the paper are present in the paper and/or the Supporting Information.

## ACKNOWLEDGMENTS

This work is supported by funding from the Cancer Prevention and Research Institute of Texas (CPRIT) award RR220008 to G.H.Z. and from the Welch Foundation Catalyst Center for Advanced Bioactive Materials Crystallization (Award V-E-0001) to G.H.Z. and A.K. The authors thank the computational resources provided by the Hewlett-Packard Enterprise Data Science Institute at the University of Houston. A.A. acknowledges the support from the Houston Endowment Fellowship. The authors thank Prof. Jens Smiatek of the University of Stuttgart for providing the codes to generate the initial configuration and topology files for PDDA polymer.

## Supporting Information Text

### SI. 1 Methods

In this work, we performed explicit-water CG MD simulations of an aqueous solution of polycationic PDDA and ATP molecules in the explicit presence of counterions as in the experimental solutions (1). We chose this model system to study polyelectrolyte coacervation phenomena because of the availability of experimental data (1, 2), where the formation of well-stable coacervates/condensates have been studied in detail.

In order to ensure adequate statistical significance, proper uncertainty quantification, and reproducibility of our findings, we performed 8 independent simulations starting from randomly monodispersed mixtures of PDDA and ATP. We calculated the standard deviation of the sample mean as an error estimate and presented them as error bars in plots whenever appropriate. We calculated the errors via the following expression, 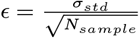 where *N*_*sample*_ is 8 (both for supernatant and DI water) and *σ*_*std*_ is the standard deviation, calculated as 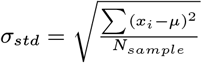 where the sum runs over *N*. *x* is the value of the corresponding observable for the given sample, and *μ* is the sample mean.

#### (a) Modeling and Force Field

We modeled the PDDA(3) and ATP(4) molecules with MARTINI 2.0 CG model in explicit water and ions. We fixed the length of each PDDA molecule as 50 monomers and each monomer has a quarternary amine group with a permanent charge of +*e*, i.e., each PDDA molecule has a total charge of +50*e*. We adjusted the protonation state of a single ATP molecule as − 4*e*, in accordance with its nearly complete dissociation behavior at pH 7 (5). The schematic CG representation implemented in this work has been provided in Figure S1. Our systems consist of 4 PDDA chains (50 monomers each) and 50 ATP molecules. This selection is made to attain charge matching between the positively and negatively charged polyelectrolytes and 20 mM ionic concentration (i.e., 20 mM PDDA monomer concentration and 5 mM ATP concentration) as examined in a recent experimental study (1). We initialized our systems as randomly dispersed PDDA and ATP molecules within a cubical box of ∼ 25 nm to achieve 20 mM ionic concentration. We also added 200 positively charged and 200 negatively charged small ion beads to mimic solution conditions (i.e., supernatant). For positively and negatively charged small ions, we used Na^+^ and Cl^−^ ion beads, respectively. We solvated the box with non-polarizable MARTINI 2.0 water (6). We refer to these simulations as the PECCs in concentrated “supernatant” conditions. We simulated each independent supernatant simulation (8 total) for at least 7 *μ*s.

We also simulated deionized (DI) water conditions by transferring the coacervates formed in supernatant conditions to pure water. In order to initialize the DI water simulations, we first isolated the coacervates formed in supernatant conditions including the PDDA, ATP, water, and ion beads enclosed in the coacervates. We described the coacervate identification protocol in *Cluster Analysis* subsection. We included all the ions and water beads at a distance smaller than 1 nm to any PDDA or ATP beads and then solvated these coacervates in pure water. We referred to the simulations as the PECCs in “DI water” conditions. See Figure S2 for a visual representation of a transferred coacervate. For each supernatant system, we have one corresponding DI water system. We refer to a given pair of supernatant and corresponding DI water system as one ‘set’, so we have a total of 8 sets. We started each independent DI water simulation (8 total) from the preformed coacervate of their corresponding supernatant simulations (as in the last saved snapshot of the corresponding supernatant system). We then simulated each independent DI water simulation (8 total) for 3.5 *μ*s. We analyzed all supernatant and DI water simulations over the last 3.5 *μ*s trajectories. Both for the supernatant and DI water system, we modeled 10% of the water beads as antifreeze type (6) to prevent freezing of the MARTINI 2.0 CG water.

We note that the PDDA force field parameters used in this work were originally developed for PDDA30 chains (3) solvated in polarizable MARTINI water (7). In order to justify the compatibility of PDDA parameters with the non-polarizable water, we carried out simulations of PDDA30 solvated in the non-polarizable MARTINI 2.0 water model and compared pair correlation function and monomer-to-monomer distance distribution with the PDDA30 in polarizable water along with all-atom simulations as performed by Vogele et al. (3). These comparisons are shown in Figure S3. Although we observe PDDA30 to be more compact when solvated in the LJ water compared to that when solvated in the polarizable water, their structural properties such as radial distribution function and distribution of monomer-to-monomer distances remained qualitatively the same, lending confidence in using the force field combination that we used in this work.

#### (b) Simulation Details

All randomly dispersed systems were energy minimized using the steepest descent algorithm prior to equilibration of the system for 100 ns in NPT ensemble where the temperature was maintained constant at 298 K with the Berendsen thermostat (8) with 1 ps time constant and pressure was maintained constant at 1 bar using Berendsen barostat (8) with a time constant of 3 ps. Following the NPT equilibration, a production run of 7 microseconds (μs) was carried out in the NPT ensemble with the same parameters with a trajectory saving frequency of 1 ns. A timestep of 20 fs was implemented for integrating Newton’s equation of motion using the leapfrog algorithm (9). The run parameters were adapted from Vogele et al. (3). A cutoff distance of 1.2 nm was used both for the van der Waals interactions and short-range Coulombic interactions. Long-range electrostatic interaction was managed by implementing the Particle Mesh Ewald (PME) method (10). All simulations have been performed by using the GROMACS 2021.4 (11–13) software package. For visualization of molecular trajectories VMD 1.9.4 (14) software has been used. For analyzing the trajectories, we have used MDAnalysis 2.2.0 (15, 16) and the OVITO program (17).

#### (c) Cluster Analysis

To quantify the coacervate formation during our simulations, we employed a clustering algorithm based on a minimum distance criterion as in one of our recent works (18). Here, we considered any ATP or PDDA in the same cluster if the minimum distance (r_*cut*_) between any PDDA or ATP beads is less than or equal to 0.85 nm. We built an adjacency matrix based on this criterion and then identified the clusters based on the connected components. Further information on the clustering algorithm can be found in this work (18). We then measured the time evolution of the fraction of PDDA and ATP molecules (*f*_*cluster*_) within the largest cluster for both supernatant and DI water systems for all simulation replicas, as depicted in Figure S4 and Figure S5 respectively. When *f*_*cluster*_ was found to reach and remain at 1.0 without any deviation, we considered the cluster to be well-formed and stable; and referred to that cluster as the coacervate.

For all the independent simulations corresponding to the supernatant system, we ensemble averaged the individual samples over the period of the last 3.5 μs (from 3.5 μs to 7 μs), whereas for DI water independent simulations, the ensemble averaging time slab was considered from 0 μs to 3.5 μs.

#### (d) Asphericity index

We also calculated the time evolution of the asphericity index (*A*_*S*_) of the largest cluster formed in both ‘supernatant’ and ‘DI water’ systems to examine the shape of the clusters. This asphericity index (*A*_*S*_) was calculated following the expression (19),

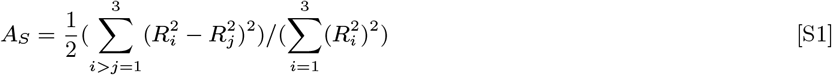

where 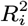 are the principal radii of gyration of the cluster represented by all of its PDDA and ATP beads. *A*_*S*_ changes between 0 and 1, where *A*_*S*_ = 0 for a perfectly spherical object and *A*_*S*_ = 1 for a rodlike object.

#### (e) Ion residence time correlation function 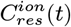

To understand the quantitative differences between ions residing in different regions of the coacervates, namely the ‘core’ and ‘surface’, we calculated the lifetimes of ions residing in those regions. This residence time was calculated from residence time correlation function 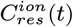via the following equation (20, 21):

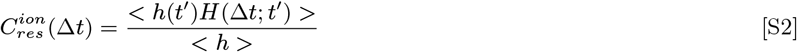

where h(*t*^*’*^) takes a value of 1 when a particle is residing in the targeted region at time *t*^*’*^ and 0 when ‘t ‘s not. *H*(Δ*t*; *t*^*’*^) takes a value of 1 if the particle is continuously residing in that region (‘core’ for residence time of core ions or ‘surface’ for residence time of surface ions) for the timespan of Δ*t* starting from time *t*^*’*^; *H*(Δ*t*; *t*^*’*^) equates to 0 if the particle goes out of the region at any given point of time within time *t*^*’*^ to *t*^*’*^ + Δ*t*. That is,

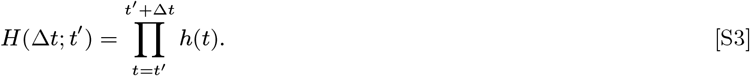

and *< h >* denotes ensemble averaged *h*(*t*^*’*^).

In this analysis, we need an explicit description of the ‘core’ and ‘surface’ of the coacervate (as well as the rest of the simulation box, which is referred to as ‘bulk’). Upon visual inspection of the coacervates (e.g., snapshots in Figure S4 and Figure S5), we observed that their shapes tend to deviate from a sphere, posing a challenge in accurately defining the surface region of the clusters. To define the surface region, we developed the following prescription:

1. The coacervate was projected on three mutually perpendicular principal planes defined by the three (dynamic) principal axes of the cluster (see Figure S6).
2. Subsequently, these projections on three mutually perpendicular planes were plotted together on a single graph using distinct colors, and the common rectangular area of overlapping projections was identified and marked.
3. We observed that the common rectangular area encloses between sets of two horizontal (at y=-2 nm, y =2 nm) and two vertical red lines (at x=-2 nm, x =2 nm). We then considered the overlapping volume in three dimensions as a sphere of radius (*r*_*inner*_ = 2.8 nm), i.e., half diagonal of this rectangular area. We consider this volume as the ‘core’ (see Figure S6).
4. Finally, the area between the inner radius (*r*_*inner*_ = 2.8 nm) and outer radius (*r*_*outer*_ = 5 nm) was considered as the interfacial region. It should be noted that *r*_*out*_=5nm is extracted from the density profile plot of PDDA and ATP beads mentioned in Figure 2A in the main text, where 5 nm was found to be the maximum radial distance from the center where PDDA or ATP beads were found. We employed this protocol for defining the ‘core’ and ‘surface’ regions for all eight sample simulations.

We presented the projections with this prescription in Figure S6 for all independent simulations in the supernatant. By analyzing these projections, we developed the following for defining the ‘core’, ‘surface’, and ‘bulk’ regions: 1. core region: the spherical region of radius *r*_*inner*_ =2.8 nm centered at the center of mass (COM) of the coacervate, 2. surface region: the spherical shell within radii *r*_*inner*_ =2.8 nm and *r*_*outer*_ =5 nm 3. bulk region: region outside the radial distance *r*_*outer*_ =5 nm from the COM of the coacervates.

## Supporting Figures

**Fig. S1.**
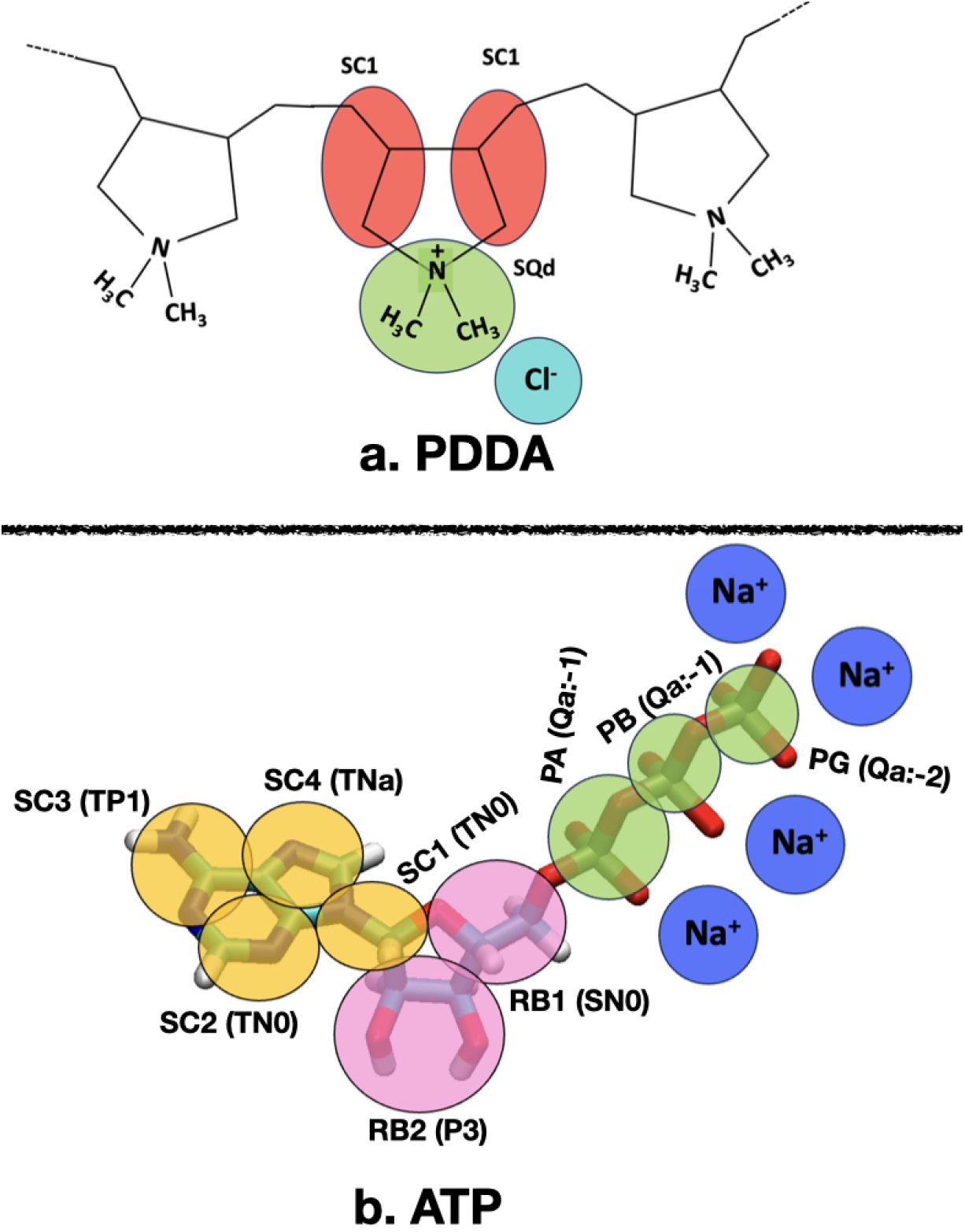
Schematic representation of atomistic to coarse-grained (CG) mapping of PDDA monomer, ATP molecule implemented in this work. The transparent colored circles represent the CG beads. A more detailed description of coarse-grained description can be found in earlier studies (3, 4).

**Fig. S2.**
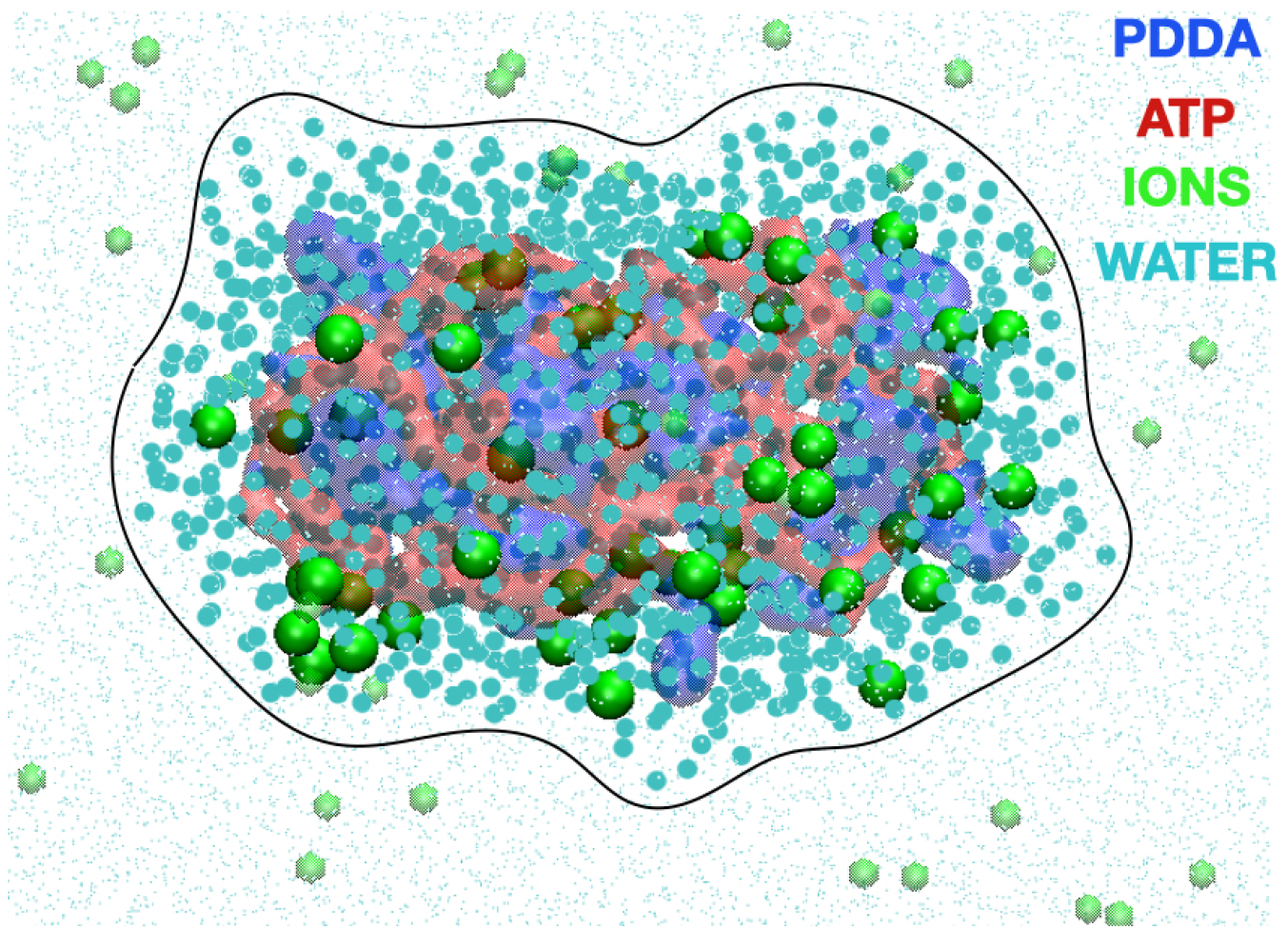
A snapshot of PDDA-ATP coacervate with associated ions and water beads just before transferring it to DI water. The following color scheme is used here: PDDA molecules are red (transparent), ATP blue (transparent), selected ions are green, and water beads retained in the coacervates are cyan. Molecules within the curved boundary (a curved surface in 3D) are the ions and waters at a distance smaller than 1 nm to any PDDA or ATP beads. Ions and waters that are discarded while transferring are shown in transparent green and cyan color with smaller bead sizes.

**Fig. S3.**
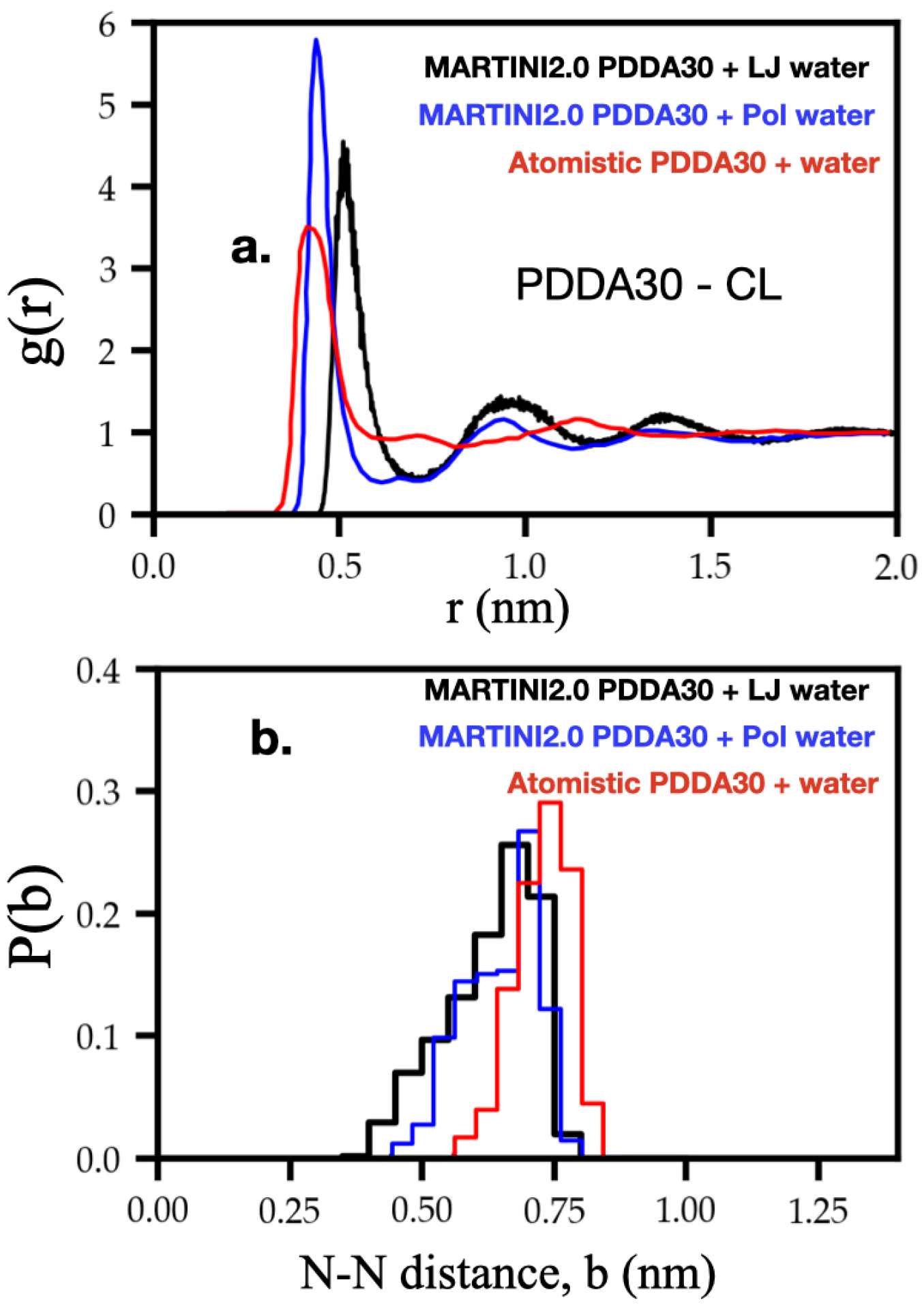
Comparison between MARTINI 2.0 PDDA30 in polarizable CG water(3), PDDA30 in LJ CG water and atomistic PDDA with atomistic water(22). In this work, we simulated PDDA30 in LJ water and compared them with existing results. (a) Radial distribution function between the charged group of PDDA30 and chloride ions. (b) Distribution of PDDA30 intramolecular monomer-to-monomer distances (i.e. distance between subsequent SQd-SQd beads for the MARTINI model and distance between N atoms for the atomistic model). PDDA30 in polarized water results are taken from Vogele et al. (3). Atomistic model results are taken from Qiao et al. (22).

**Fig. S4.**
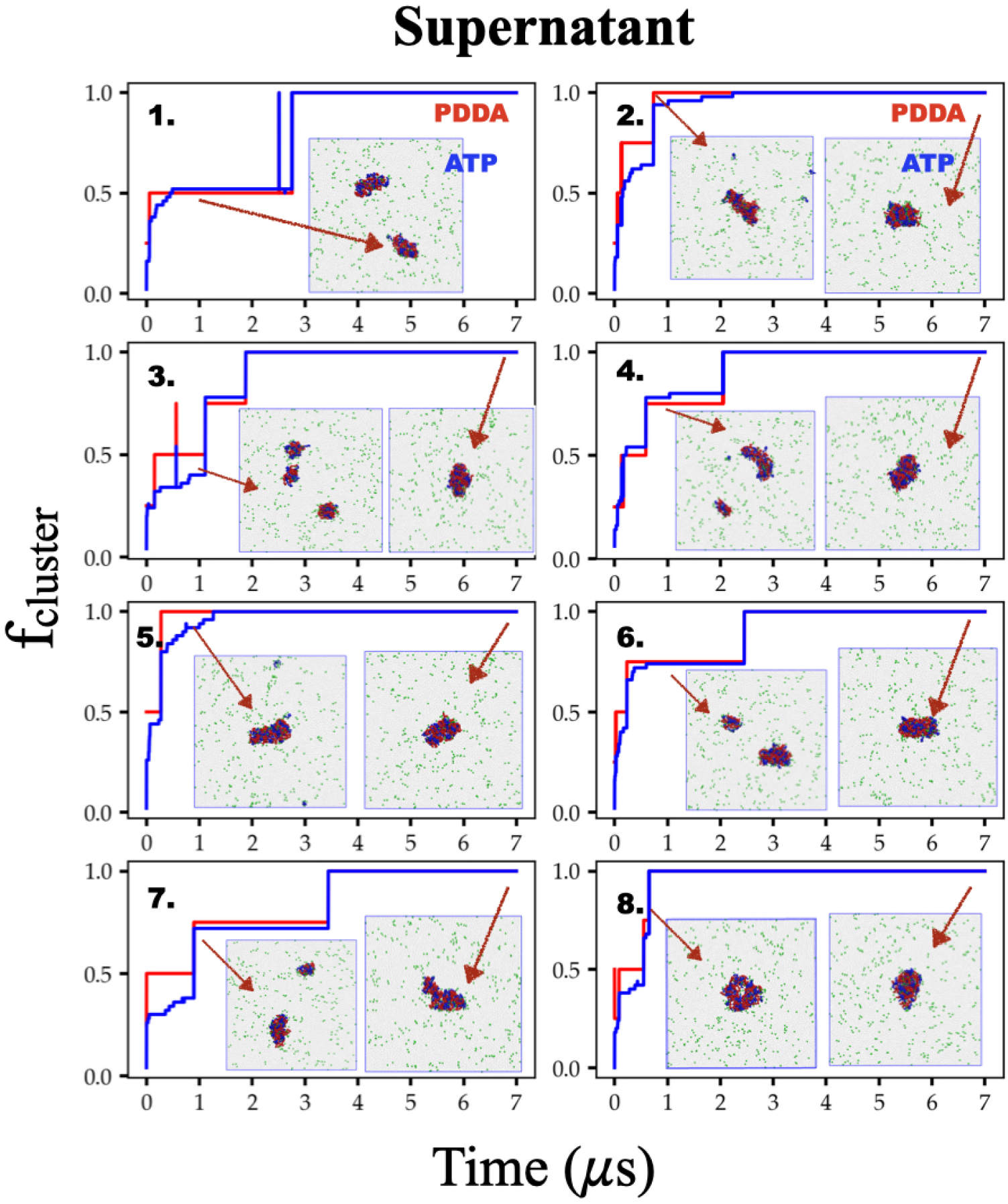
Time evolution of the fraction of PDDA and ATP molecules in the largest cluster (*f*_*cluster*_) in the ‘supernatant’ system for all the independent simulations. We consider the cluster to be stable after *f*_*cluster*_ reaches a stable value of 1.0. The minimum distance (r_*cut*_) between the beads is 0.85 nm. Snapshots provided as insets represent the cluster(s) for visualization at the simulation times indicated by arrows.

**Fig. S5.**
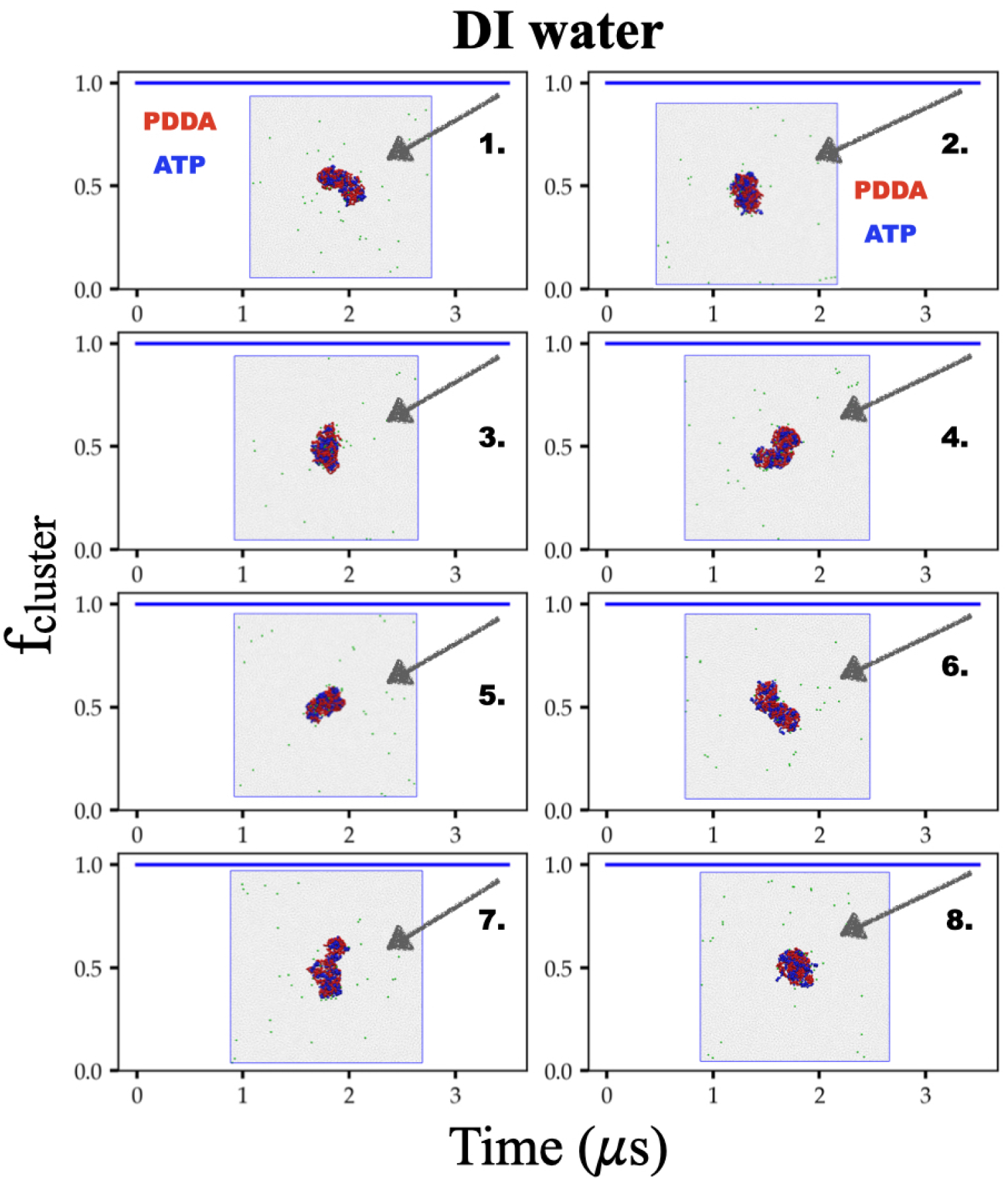
Time evolution of the fraction of PDDA and ATP molecules in the largest cluster (*f*_*cluster*_) in the ‘DI water’ system for all the independent simulations. We consider the cluster to be stable after *f*_*cluster*_ reaches a stable value of 1.0. The minimum distance (r_*cut*_) between the beads is 0.85 nm. Snapshots provided as insets represent the cluster(s) for visualization at the simulation times indicated by arrows. In ‘DI water’ clusters formed by PDDA-ATP molecules are stable against dissociation.

**Fig. S6.**
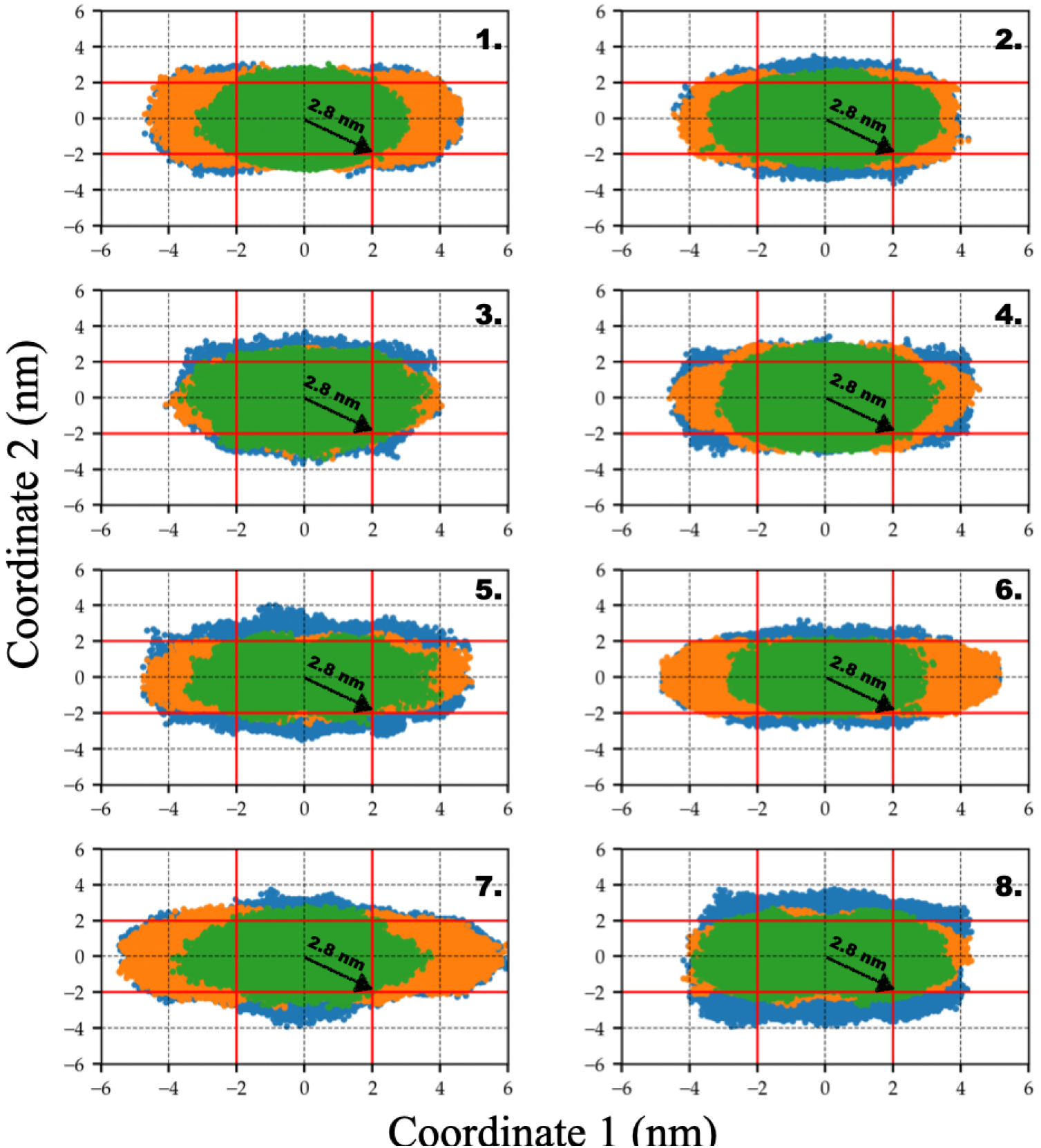
Projection of clustered PDDA and ATP beads on three principal planes made up of three dynamic principal axes of the cluster formed within the ‘supernatant’ system for multiple independent simulations to define the ‘core’ and ‘surface’ region. For all our analysis, we consider one unique prescription to define core and surface region: a. the spherical region within the radius *r*_*core*_=2.8 nm from the COM of the cluster is the ‘core’ region, b. the spherical volume within radii *r*_*core*_=2.8 nm and *r*_*surface*_=5 nm is considered as the ‘surface’ region. Here, coordinate 1 and coordinate 2 are the possible pair of principal axes of the stable cluster. Three colored regions are projections of PDDA and ATP beads on three mutually perpendicular principal planes.

**Fig. S7.**
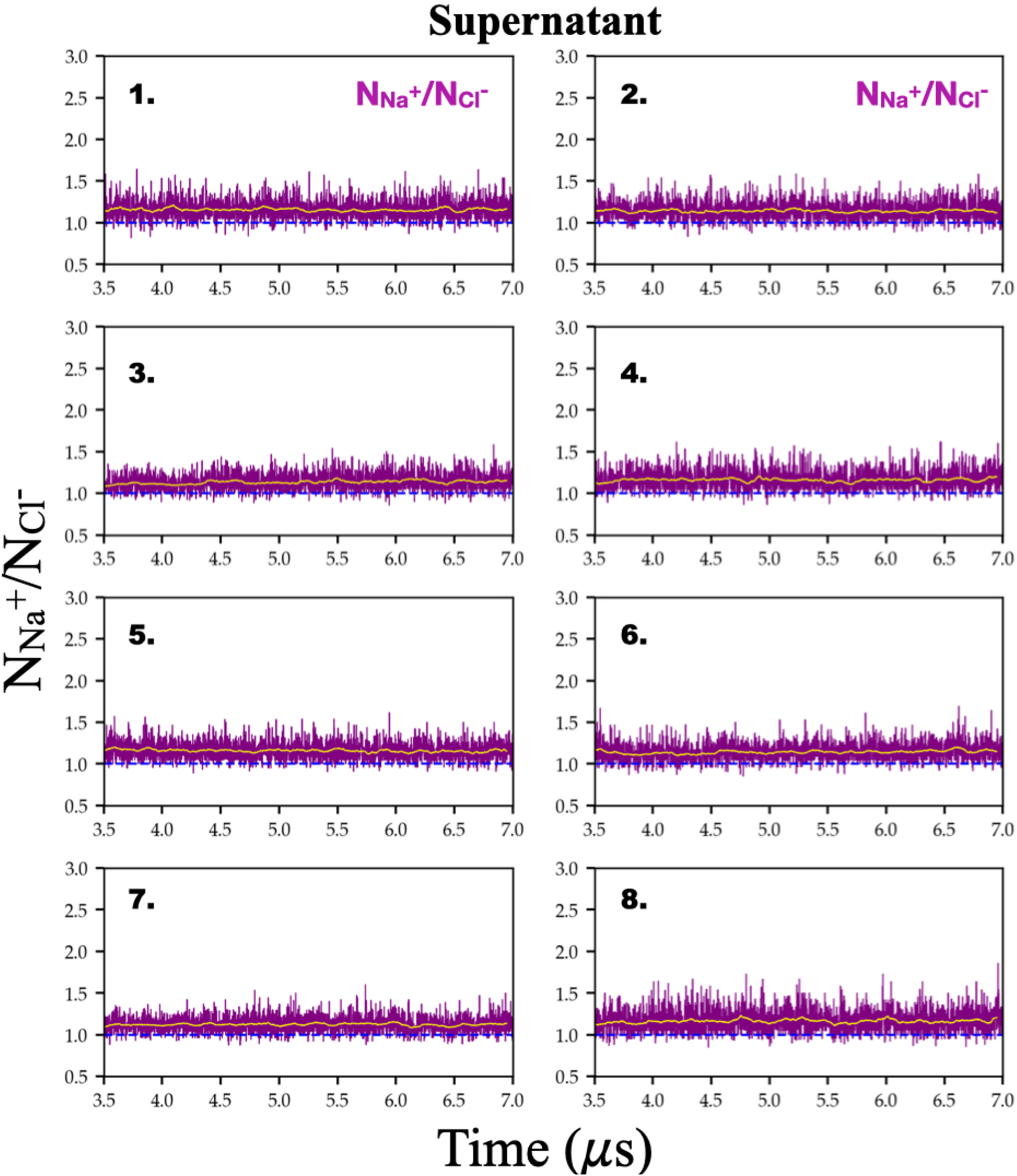
Time evolution of the ratio of the number of *Na*^+^ to *Cl*^−^ ions residing within the coacervates formed in the ‘supernatant’ system across all the independent simulations. Our well-equilibrated PDDA-ATP coacervates are always positively charged.

**Fig. S8.**
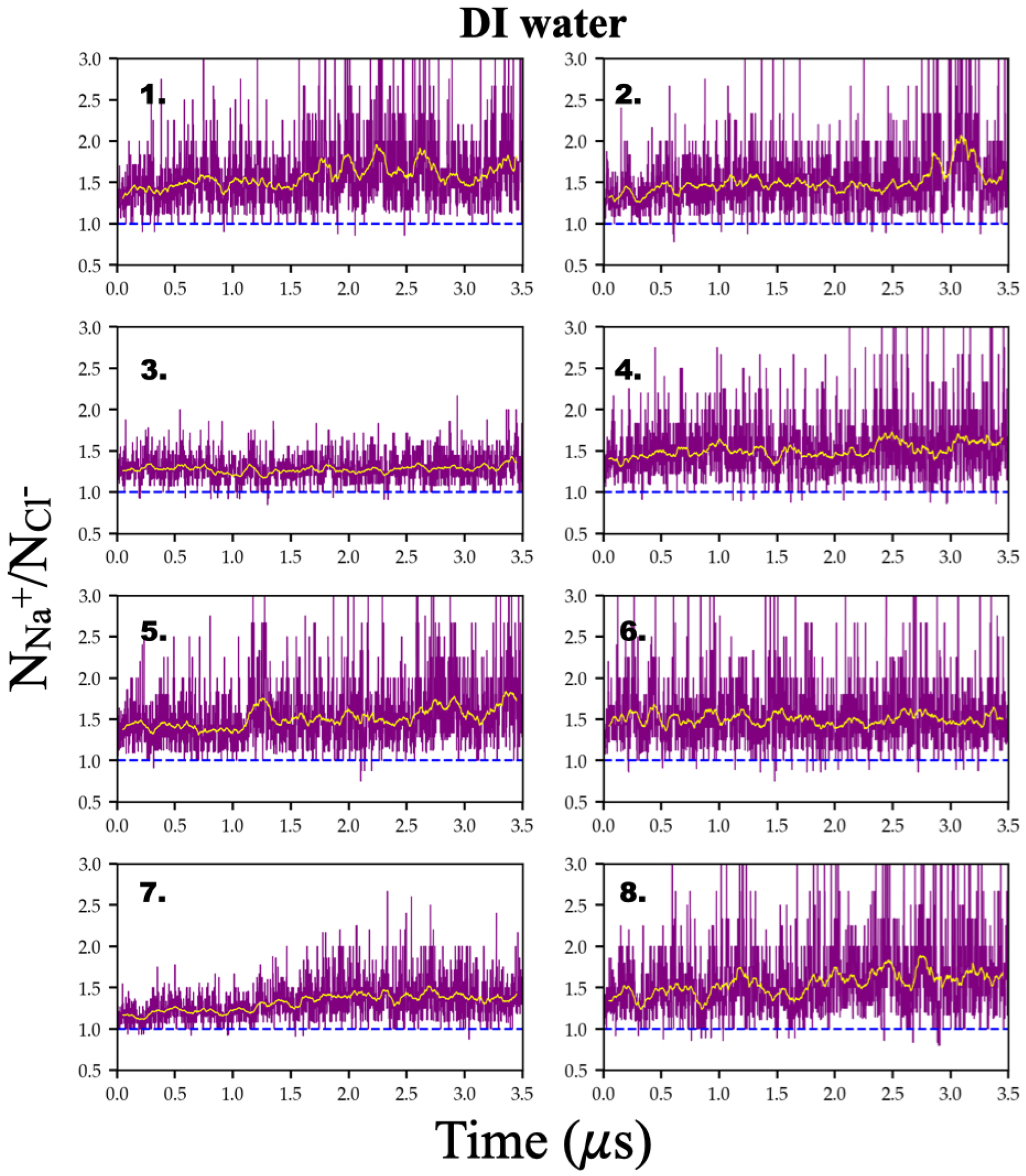
Time evolution of the ratio of the number of *Na*^+^ to that of *Cl*^−^ ions within the coacervates in ‘DI water’ systems across all the independent simulations. When well-equilibrated PDDA-ATP coacervates are transferred to ‘DI water’, they always become more positively charged. The order of panels in this figure and that in Figure S7 follow the same identity of a given set to ensure one-on-one comparison.

**Fig. S9.**
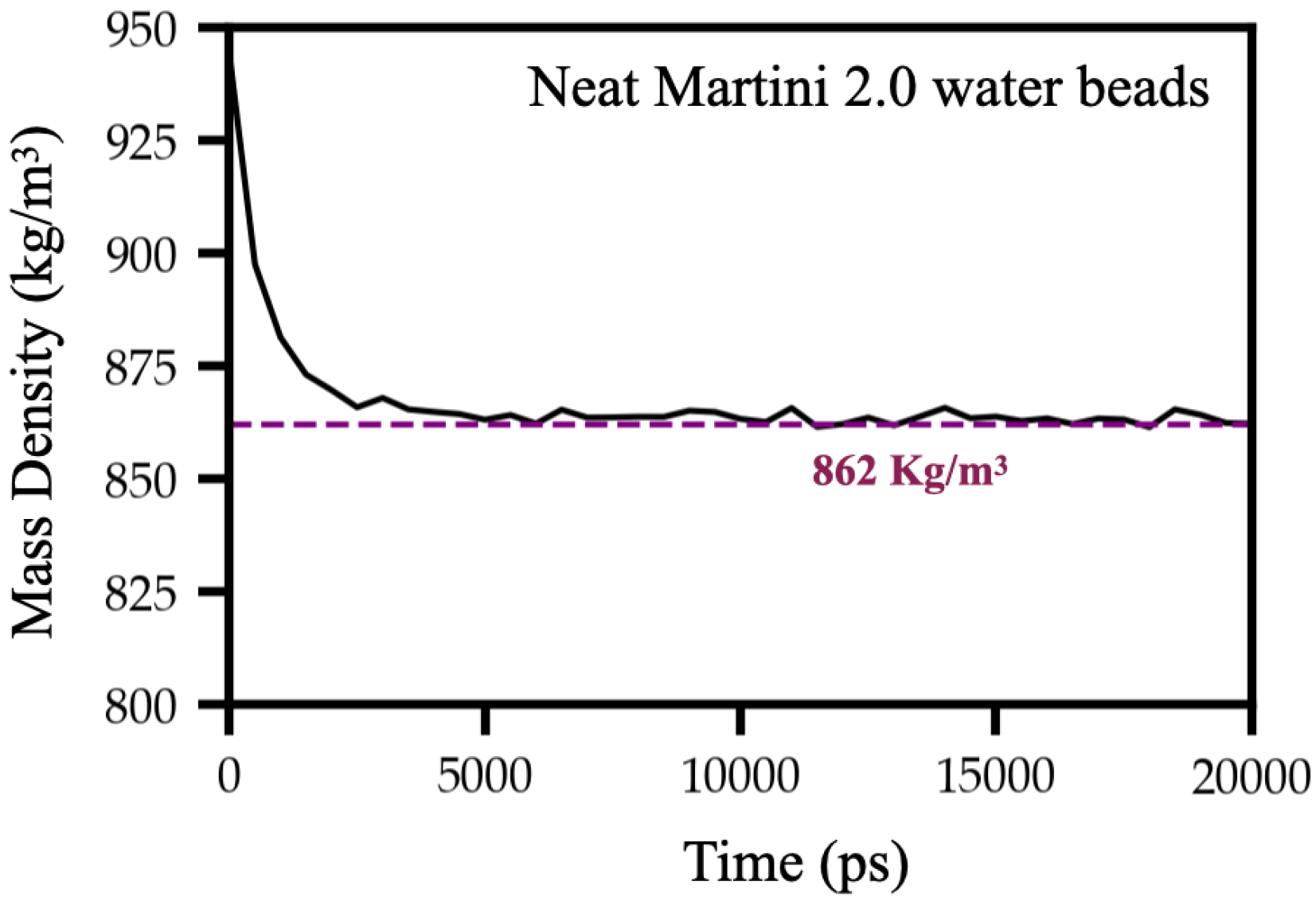
Time evolution of the density of pure MARTINI2.0 water at T=298K and P=1 bar. Equilibrated neat MARTINI2.0 water density has been found to converge at 862 kg/m^3^.

**Fig. S10.**
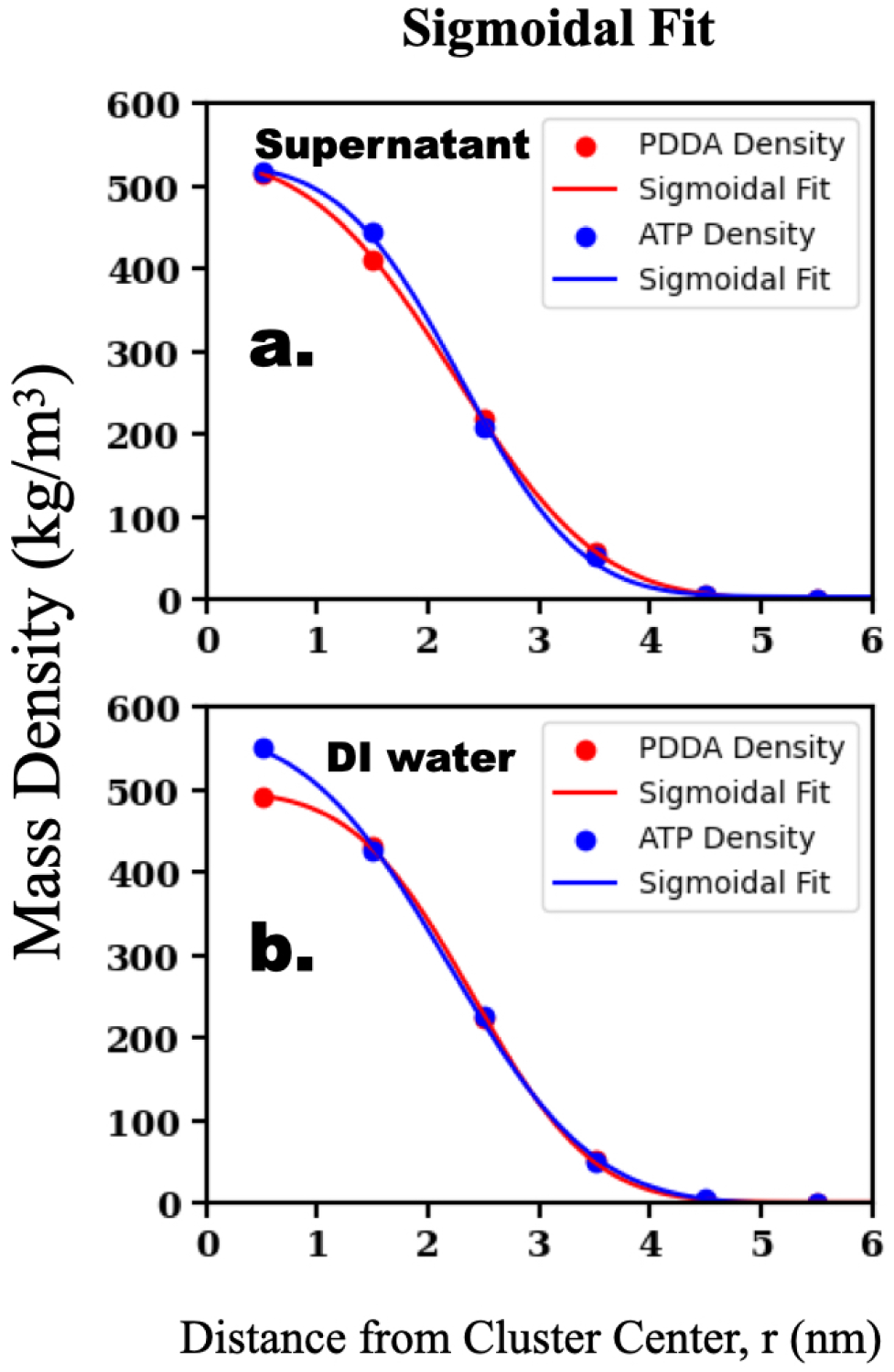
Sigmoidal fit of the density of PDDA and ATP molecules in ‘supernatant’ (a) and in ‘DI water’ (b). Sigmoidal fit function 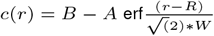 is used following earlier studies. (18, 23) For relatively larger clusters, the polymer density in coacervates (that is, the dense phase density, *C*_*dense*_) can be estimated as (B+A) and the polymer density in the dilute phase (*C*_*dilute*_) can be estimated as (B-A). For PDDA, we estimated the *C*_*dense*_ in ‘supernatant’ as 539.4 kg/m^3^, and that in ‘DI water’ as 511.3 kg/m^3^ using sigmoidal fits. For ATP, the *C*_*dense*_ in ‘supernatant’ is 535.9 kg/m^3^ and that in ‘DI water’ is 565.9 kg/m^3^. In both ‘supernatant’ and ‘DI water’, the dilute phase density (*C*_*dilute*_) is practically zero.

**Fig. S11.**
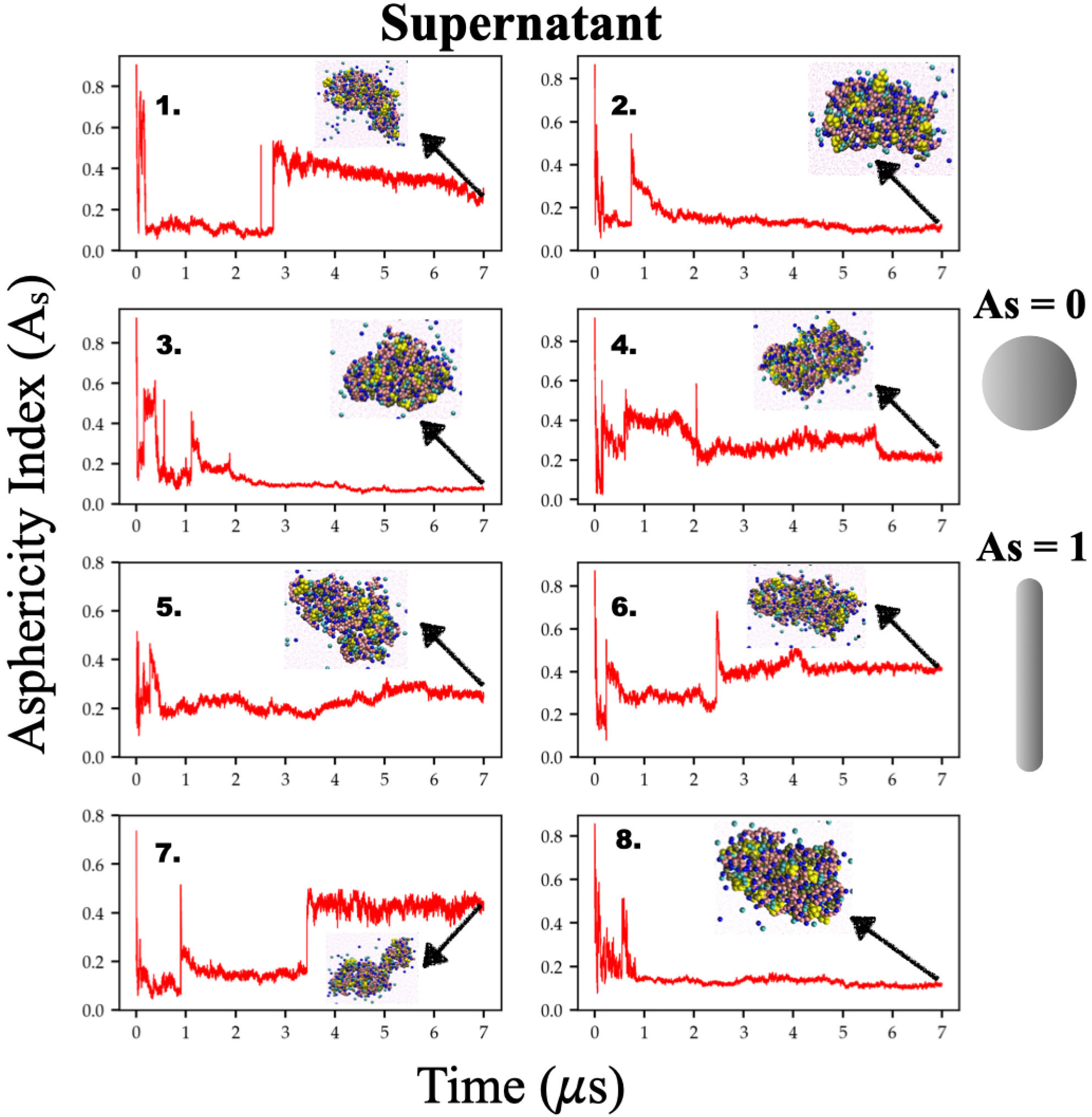
Time evolution of asphericity index (*A*_*s*_) of the largest cluster in the ‘supernatant’ system for all independent simulations. The insets display snapshots of the cluster shape captured during the final frame of the simulation (i.e. 7*μs*).

**Fig. S12.**
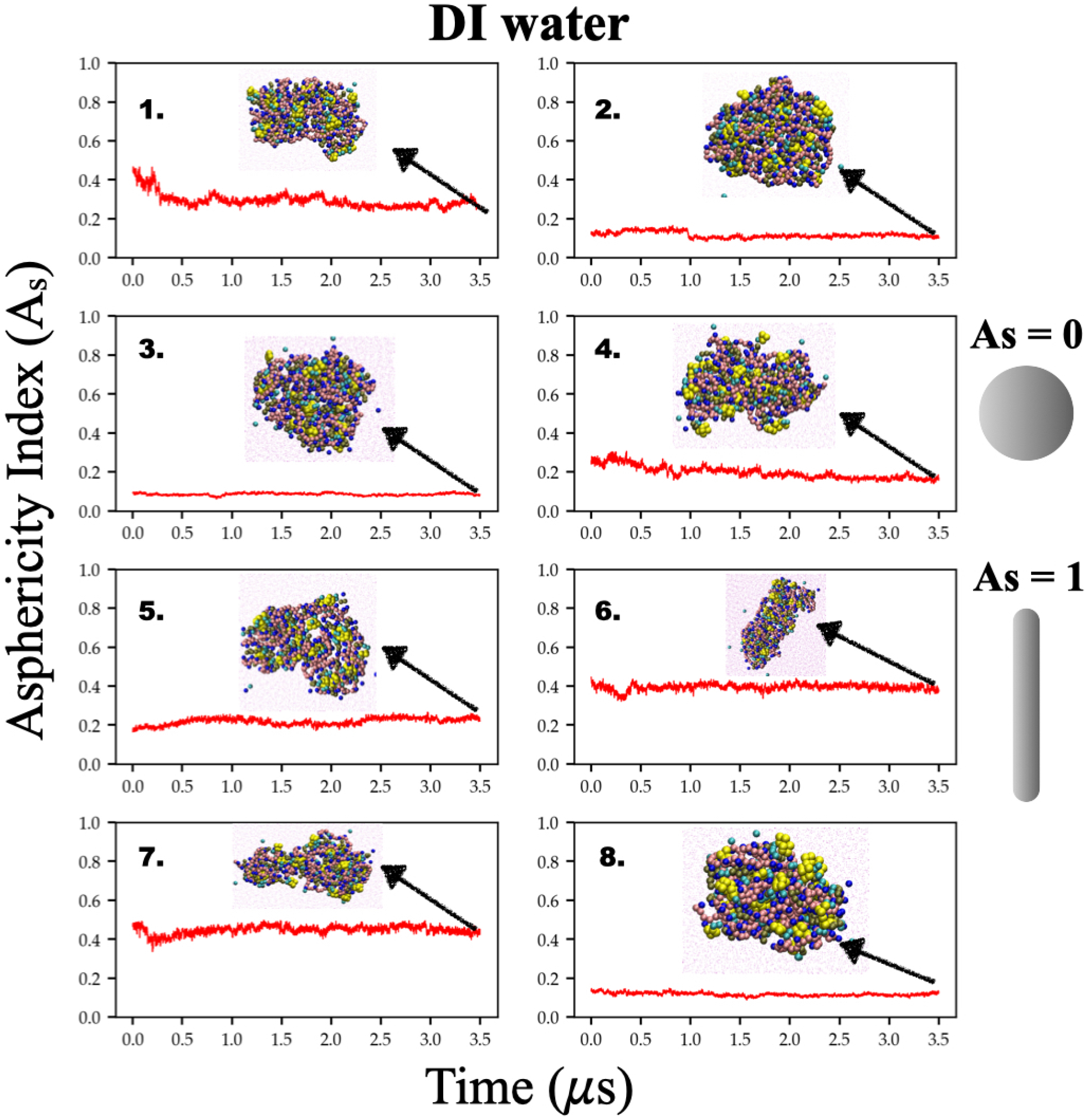
Time evolution of asphericity index (*A*_*s*_) of the largest cluster in the ‘DI water’ system for all independent simulations. The insets display snapshots of the cluster shape captured during the final frame of the simulation (i.e. 3.5*μs*). After transferring the cluster to ‘DI water’, its morphology exhibits minimal variation, characterized by a low degree of fluctuations in *A*_*s*_.

**Fig. S13.**
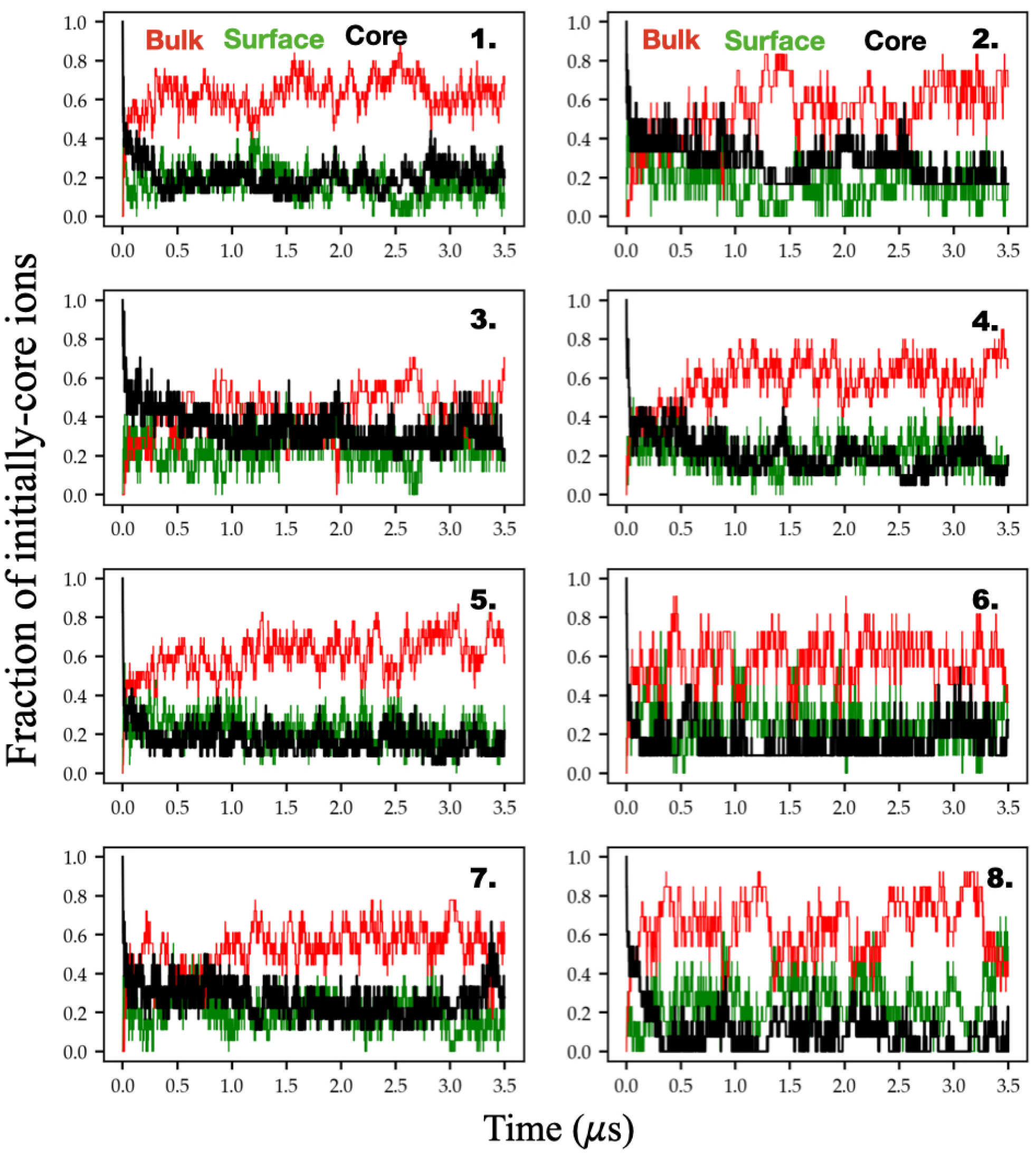
Time evolution of the fraction of initially-core ions in ‘core’, ‘surface’, and ‘bulk’ regions for all independent simulations in DI water system. Core ions are tagged at time t=0 ns (just after transferring to DI water) located inside the spherical region of radius *r*_*core*_=2.8 nm centered at the center of mass (COM) of the coacervates. After transferring to DI water 40%-60% population of the initially tagged core-ions move to the ion-depleted bulk region.

**Fig. S14.**
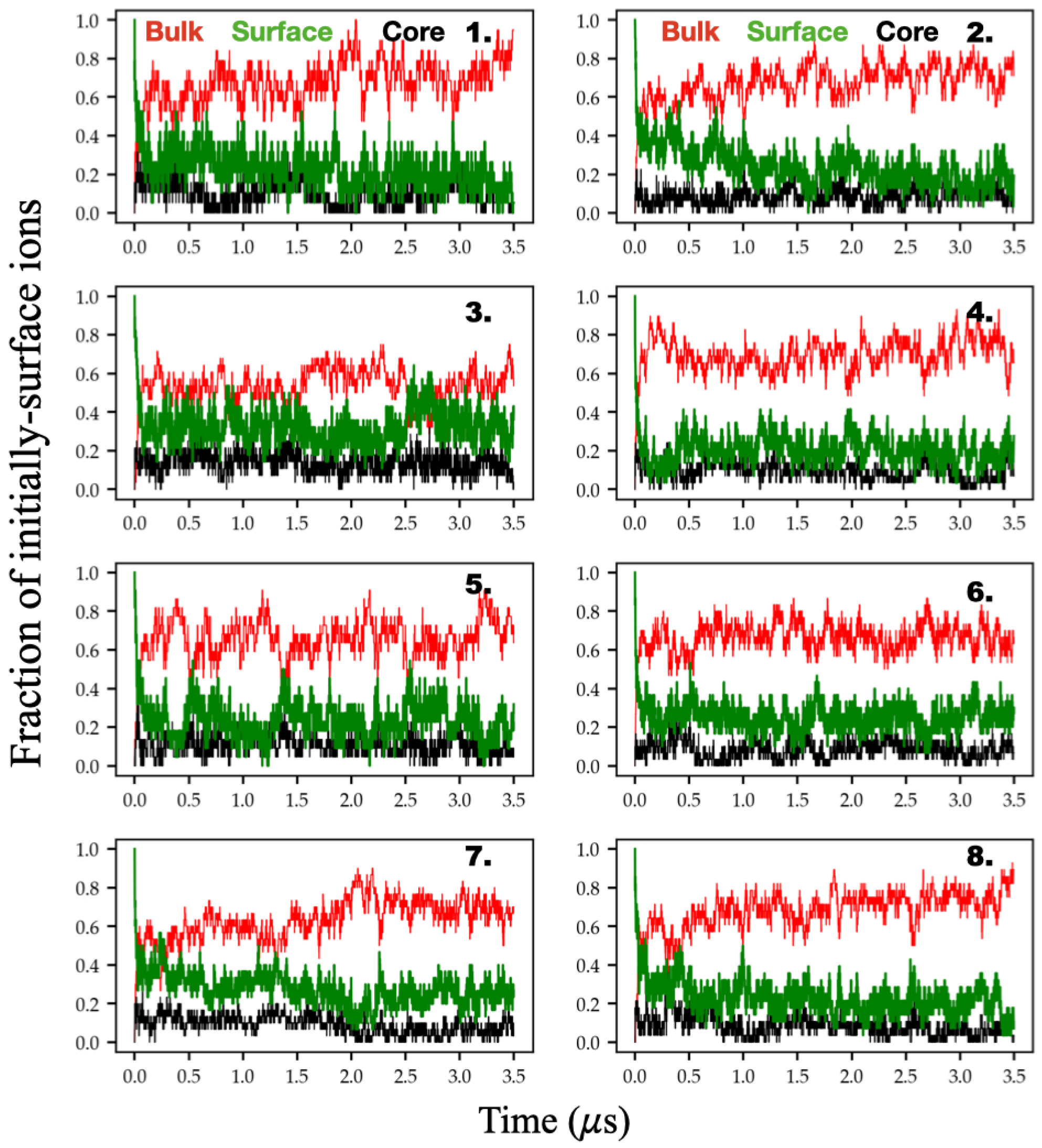
Time evolution of the fraction of initially-surface ions in ‘core’, ‘surface’, and in ‘bulk’ regions for all independent simulations in DI water system. Surface ions are tagged at time t=0 ns (just after transferring to DI water) located in the hollow sphere with inner radius *r*_*core*_=2.8 nm and outer radius *r*_*surface*_=5 nm (from the COM of the cluster). After transferring to DI water 60%-80% population of the initially tagged surface-ions move to the ion-depleted bulk region.

**Fig. S15.**
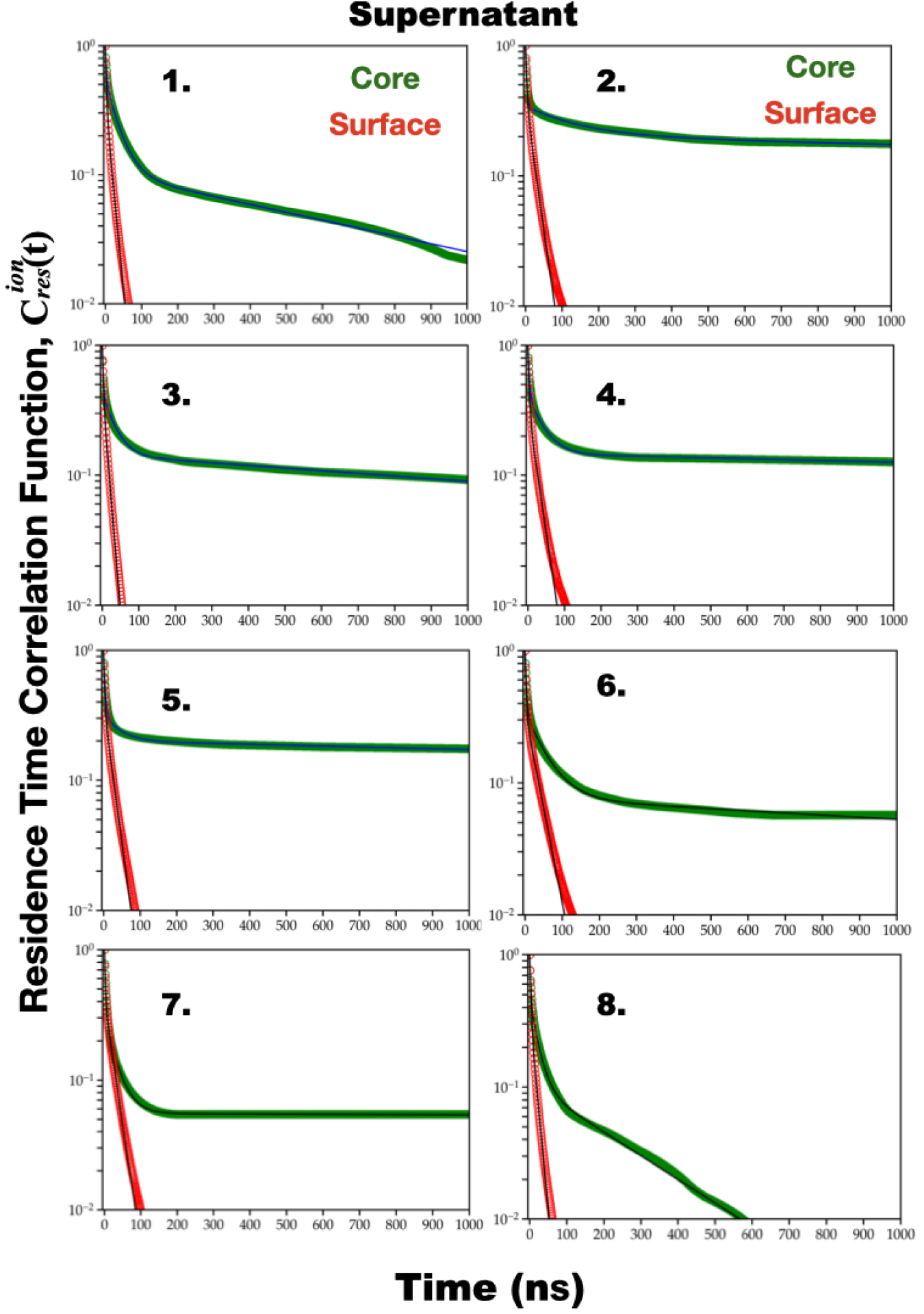
Ion residence correlation decays in the ‘surface’ and ‘core’ regions for all ions in the supernatant system across all independent simulations. Decays associated with ions residing within the ‘core’ region have been fitted via tri-exponential functions, whereas those for the ‘surface’ region are found to be adequately described by bi-exponential functions. Thin black lines are fits whose parameters are tabulated in Table S1 - S2.

**Fig. S16.**
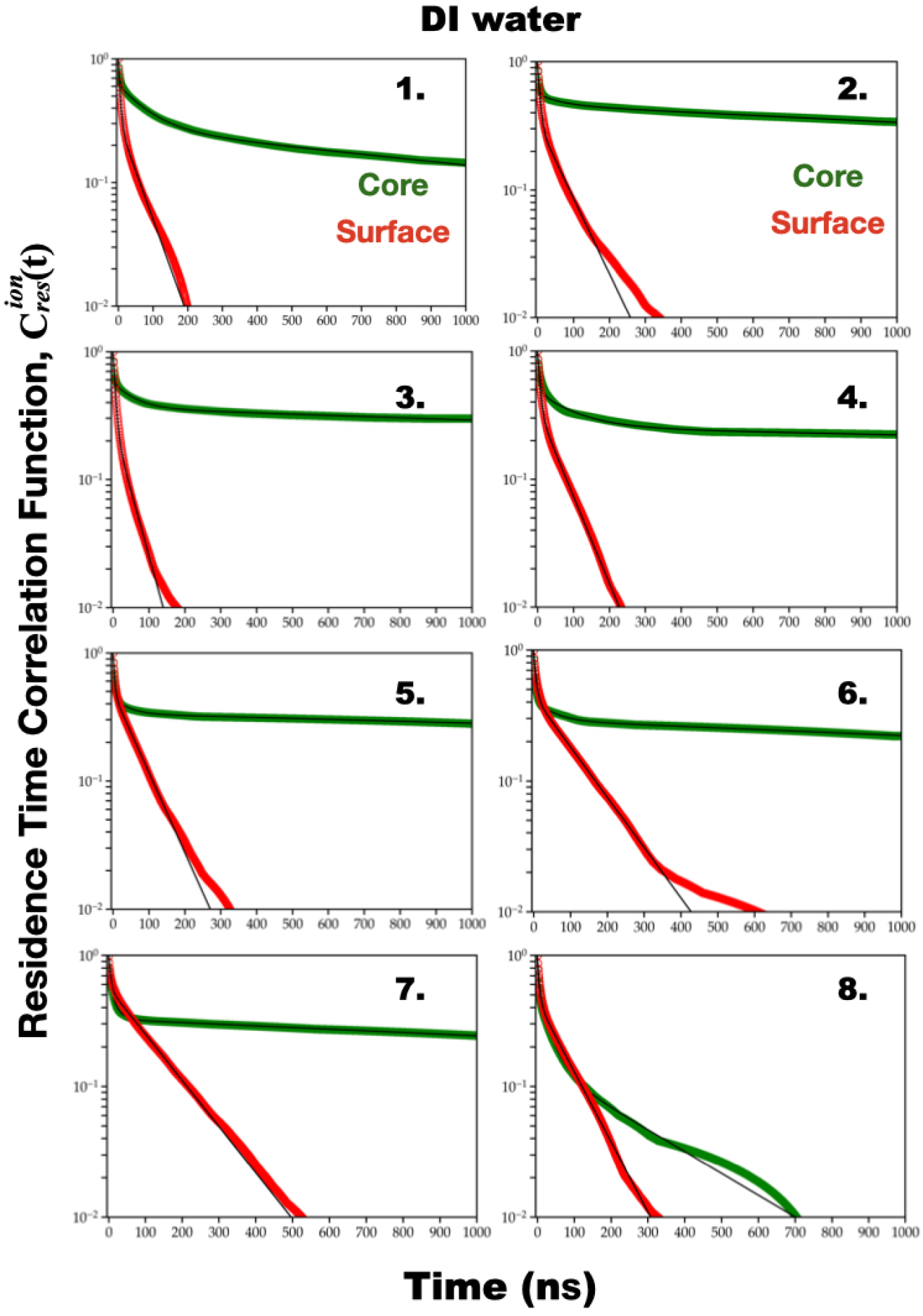
Ion residence correlation decays in the ‘surface’ and ‘core’ regions for all ions in the DI water system across all independent simulations. Decays associated with ions residing within the ‘core’ region have been fitted via tri-exponential functions, whereas those for the ‘surface’ region are found to be adequately described by bi-exponential functions. Thin black lines are fits whose parameters are tabulated in Table S3 - S4.

**Fig. S17.**
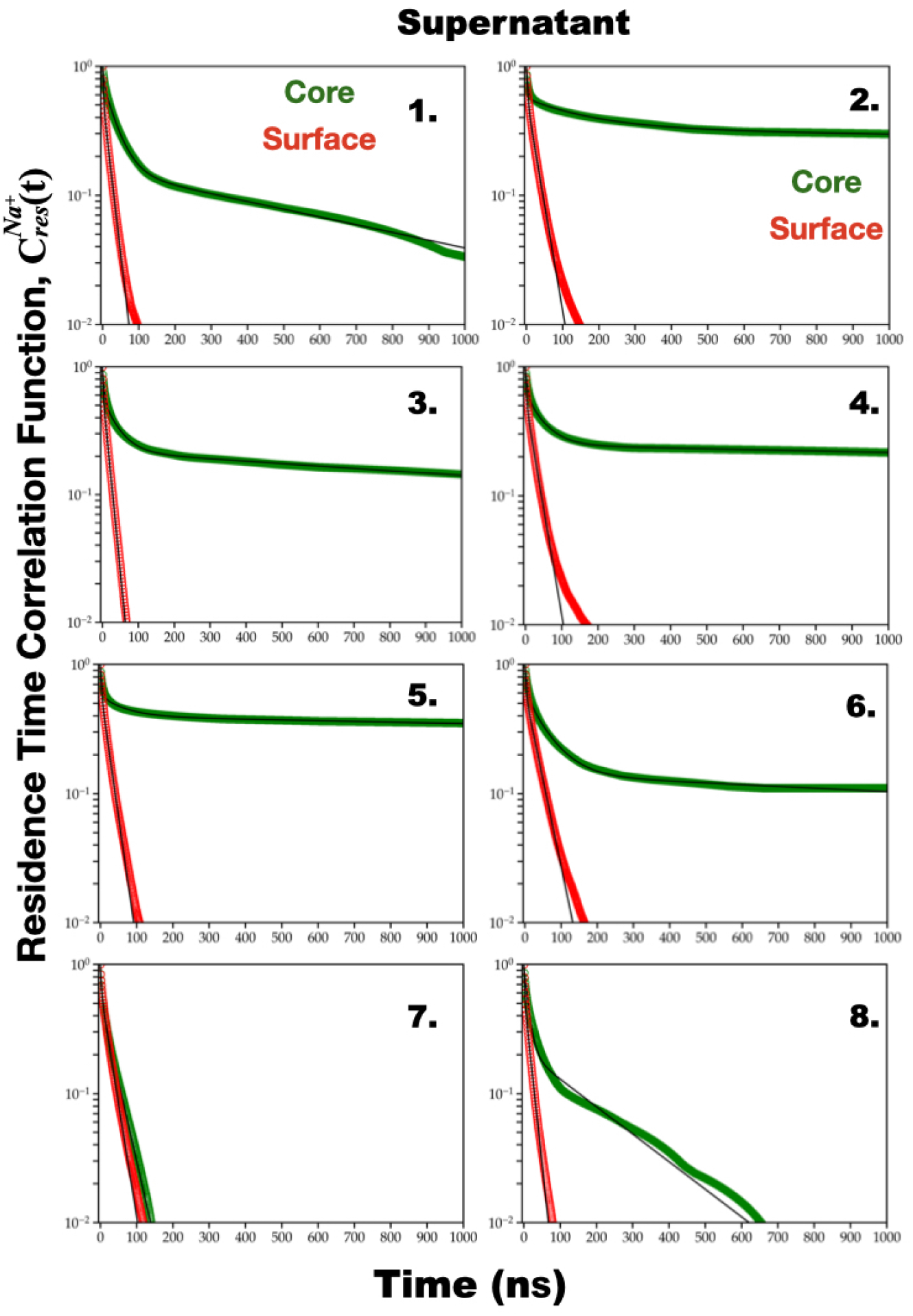
Ion residence correlation decays in the ‘surface’ and ‘core’ region for *Na*^+^ ions in the supernatant system across all independent simulations. Decays associated with *Na*^+^ ions residing within the ‘core’ region have been fitted via tri-exponential functions, whereas those for the ‘surface’ region are found to be adequately described by bi-exponential functions. Thin black lines are fits whose parameters are tabulated in Table S5 - S6.

**Fig. S18.**
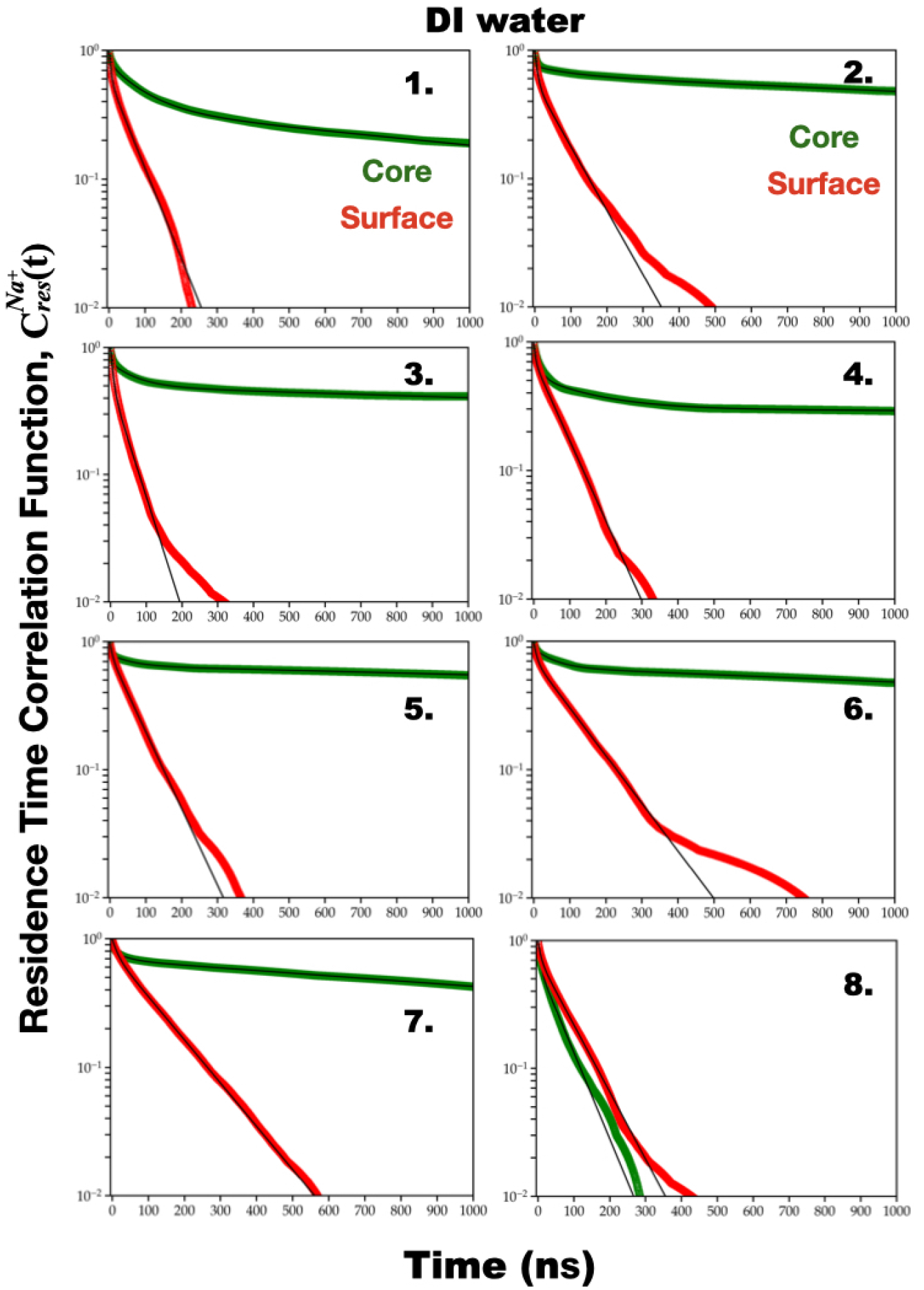
Ion residence correlation decays in the ‘surface’ and ‘core’ region for *Na*^+^ ions in DI water system across all independent simulations. Decays associated with *Na*^+^ ions residing within the ‘core’ region have been fitted via tri-exponential functions, whereas those for the ‘surface’ region are found to be adequately described by bi-exponential functions. Thin black lines are fits whose parameters are tabulated in Table S7 - S8.

**Fig. S19.**
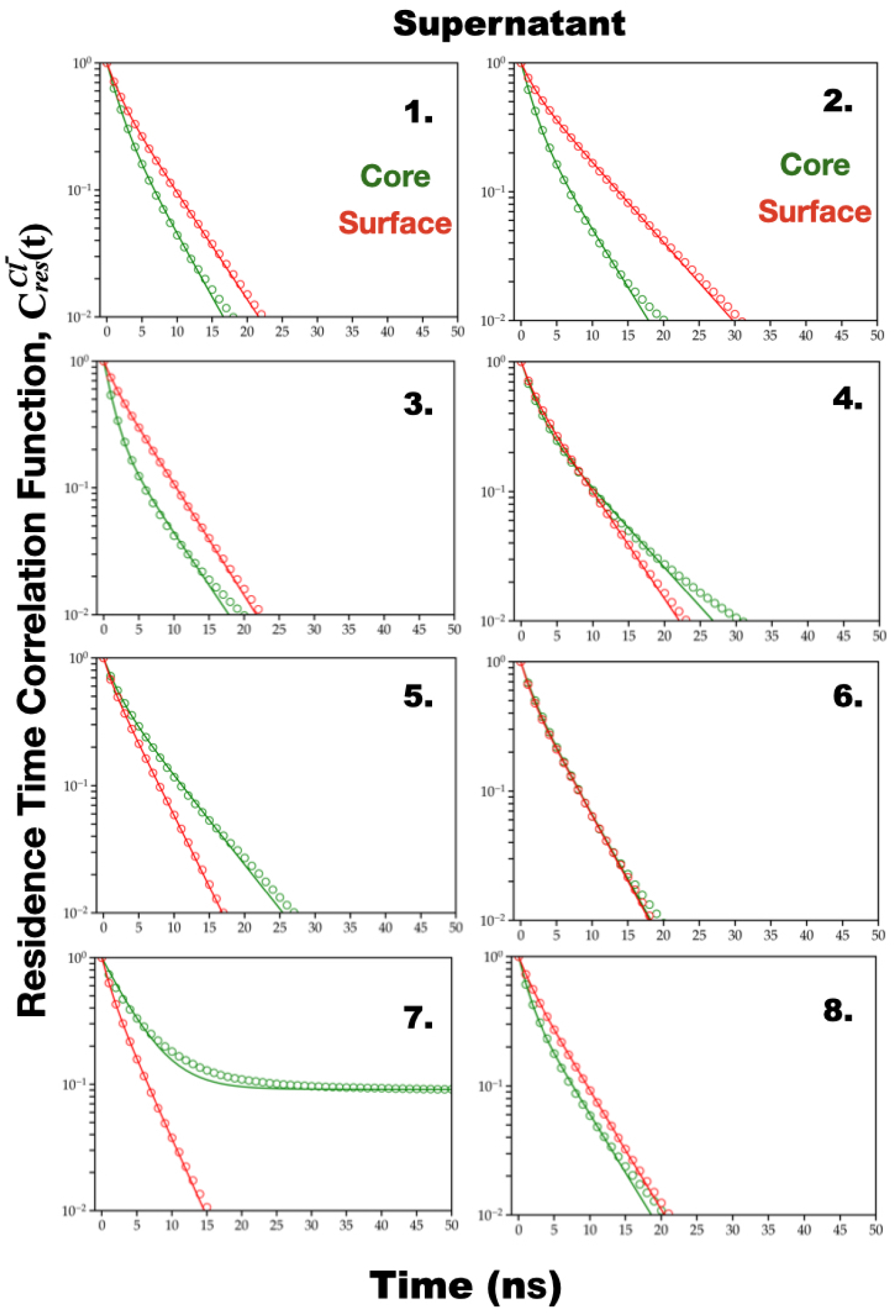
Ion residence correlation decays in the ‘surface’ and ‘core’ region for *Cl*^−^ ions in the supernatant system across all independent simulations. Most of the decays associated with *Cl*^−^ ions residing within both the ‘core’ and ‘surface’ regions are found to be adequately described by single-exponential fits. Note that the decay of *Cl*^−^ ions within the core region of independent simulation number 7 is found to fit better to a bi-exponential function. Thin black lines are fits whose parameters are tabulated in Table S9 - S10.

**Fig. S20.**
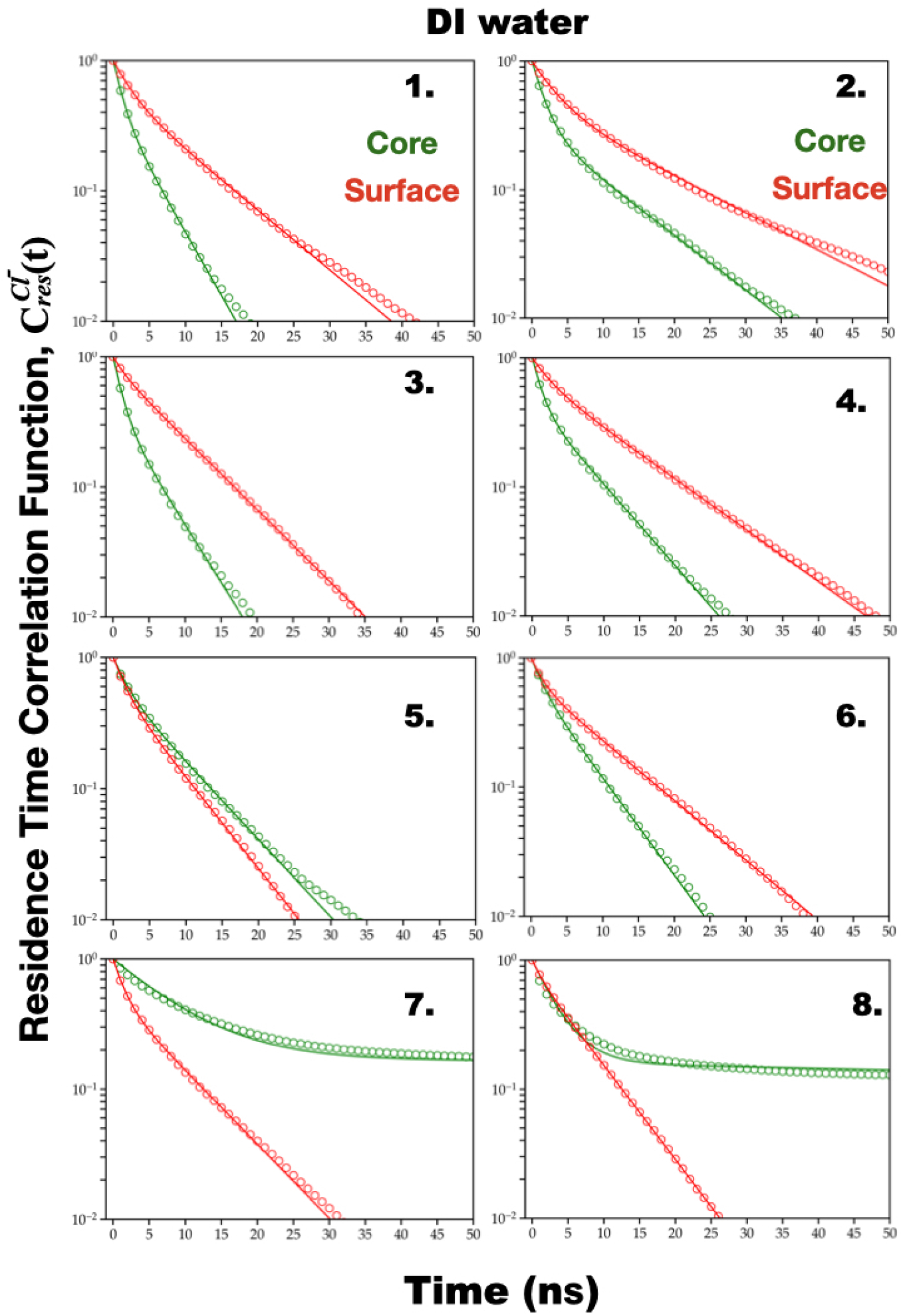
Ion residence correlation decays in the ‘surface’ and ‘core’ regions for *Cl*^−^ ions in the DI water system across all independent simulations. Most of the decays associated with *Cl*^−^ ions residing within both the ‘core’ and ‘surface’ regions are found to be adequately described by single-exponential fits. Note that the decay of *Cl*^−^ ions within the core region of independent simulation numbers 7, and 8 fit better to a bi-exponential fit function. Thin black lines are fits whose parameters are tabulated in Table S11 - S12.

**Fig. S21.**
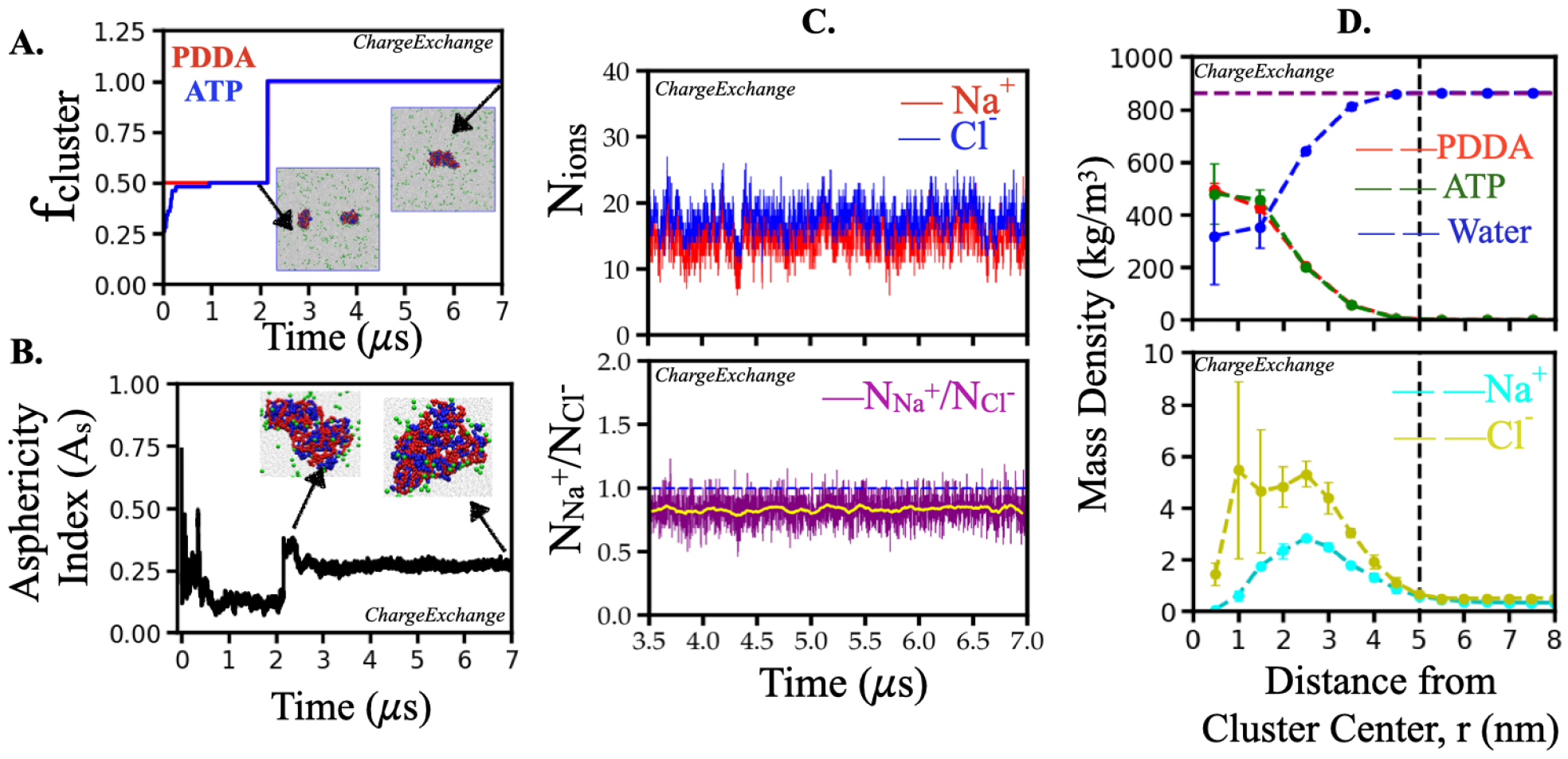
Coacervate formation, shape, size and the ionic nature of the coacervates in hypothetical ChargeExchange simulation in ‘supernatant’. A. Time evolution of the fraction of PDDA and ATP molecules in the largest cluster (*f*_*cluster*_) for a selected set. The snapshots in the insets are the simulation box at indicated simulation times where PDDA, ATP, and small ions beads are colored red, blue, and green, respectively. B. Time evolution of asphericity index of the largest cluster for a selected set. Attached snapshots in the inset are the clusters at indicated simulation times where PDDA, ATP, and ion beads are colored in red, blue, and green, respectively. C. Time evolution of the number of *Na*^+^ and *Cl*^−^ ions within the coacervate for a selected set (upper panel). Time evolution of the ratio of the number of *Na*^+^ to *Cl*^−^ ions within the coacervate (lower panel). Yellow lines in the lower panel indicate a running average of the ratio of the number of *Na*^+^ to *Cl*^−^ ions, which is lower than 1 contrary to the observation in simulation of actual PDDA-ATP-water-ion simulations employing MARTINI2.0 force field. D. Mass density profiles of PDDA, ATP, water, and ions. Error bars are the standard deviation of sample means. The density profiles are averaged over two independent ChargeExchange simulations. The vertical line at 5 nm is the guiding line indicating the maximum extent of the cluster. The horizontal line at 862 kg/m^3^ refers to equilibrated MARTINI 2.0 water density at 298 K. The molecular weight of each PDDA monomer in its fully dissociated state is 126.21 g/mol and that of an ATP molecule at the protonation state of −4*e* is 503.2 g/mol. The molecular weight of water, *Na*^+^ and *Cl*^−^ ions are 18.01, 22.99, and 35.45 g/mol, respectively.

**Fig. S22.**
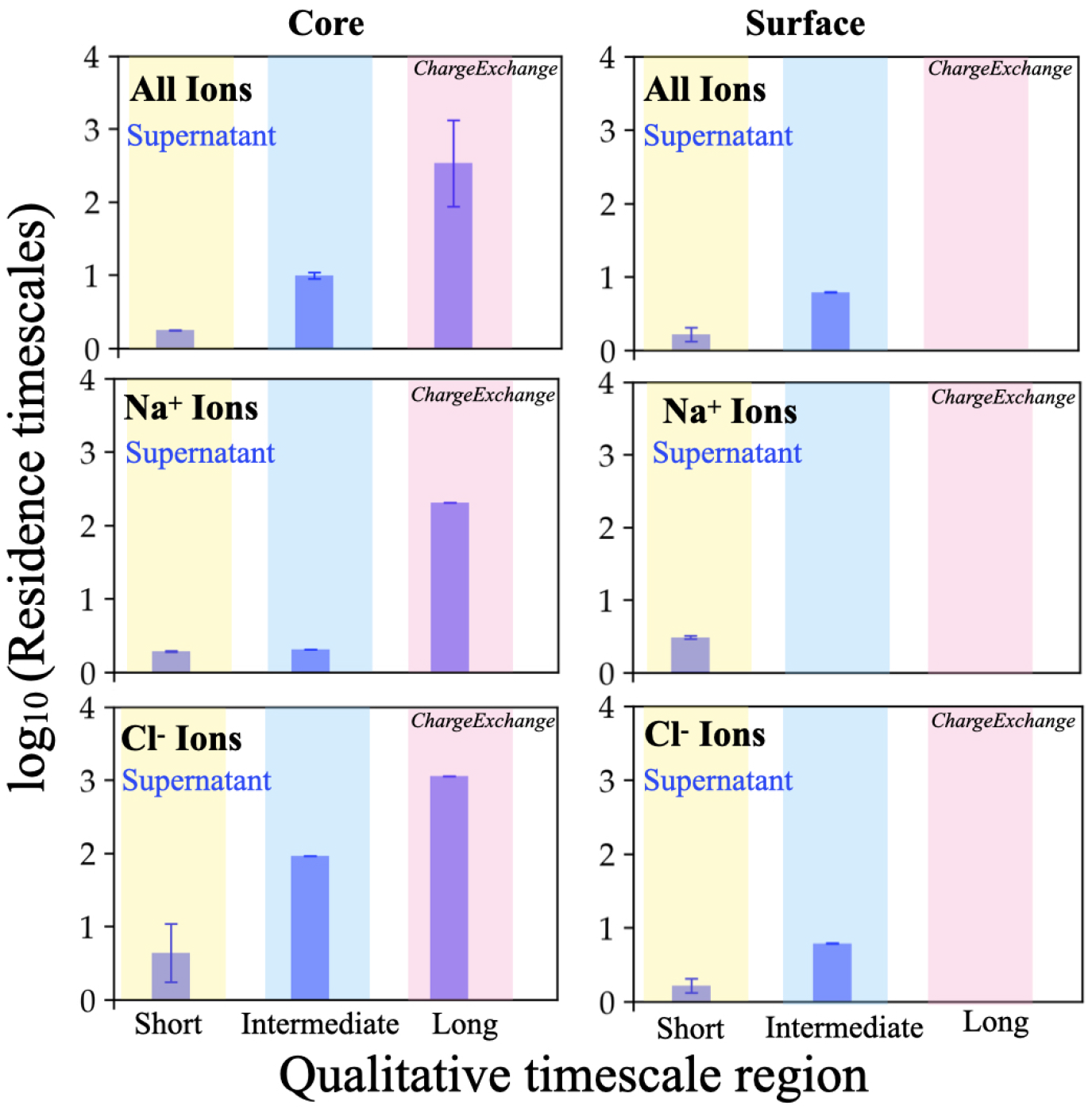
Comparison of characteristic timescales (*log*_10_(*τ*)) describing ion residence correlation decays in ‘core’ (left panel) and ‘surface’ region (right panel) of coacervates formed in hypothetical ‘ChargeExchange’ simulations in ‘supernatant’. The top panels report the timescales associated with overall ion dynamics whereas the middle and bottom panels report the same separately for *Na*^+^ ions and *Cl*^−^ ions, respectively. Average values and associated error bars (standard deviation of sample mean) are calculated over two copies of this hypothetical simulations.

## Supporting Tables

**Table S1.**
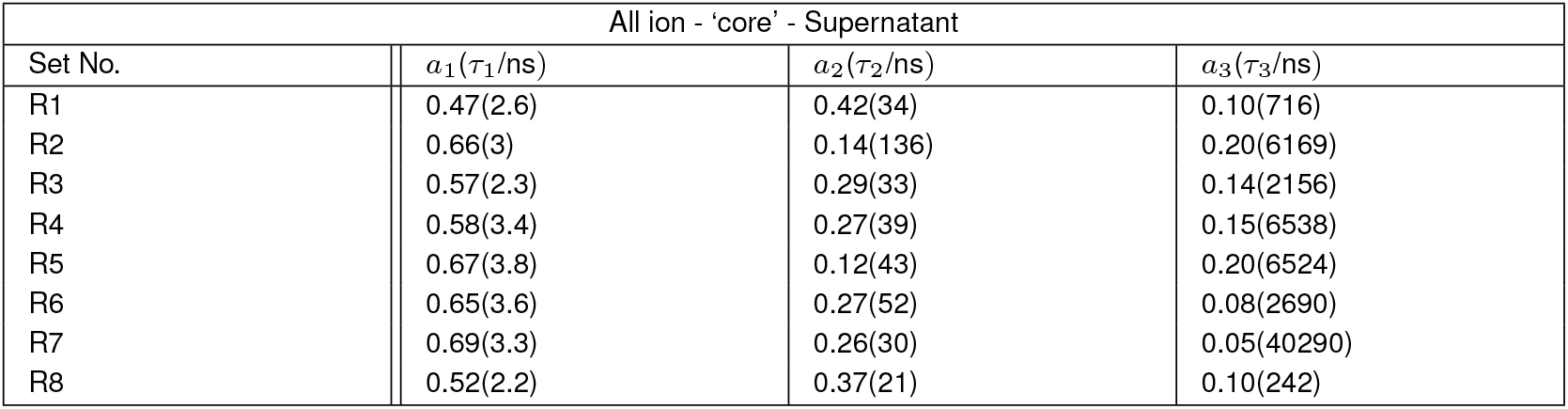
Tri-exponential fit parameters for residence time correlation function for all ions in the ‘core’ region of the coacervate in the supernatant system. Fit parameters for all eight independent simulations are presented here.

**Table S2.** Bi-exponential fit parameters for residence time correlation function for all ions in the ‘surface’ region of the coacervate in the supernatant system. Fit parameters for all eight independent simulations are presented here.

| All ion - ‘surface’ - Supernatant |  |  |
| --- | --- | --- |
| Set No. | $a_1(\tau_1/\text{ns})$ | $a_2(\tau_2/\text{ns})$ |
| R1 | 0.65(2.6) | 0.35(15) |
| R2 | 0.62(3.7) | 0.38(22) |
| R3 | 0.57(2.7) | 0.43(13) |
| R4 | 0.68(3.1) | 0.32(23) |
| R5 | 0.63(2.7) | 0.37(21) |
| R6 | 0.66(2.8) | 0.34(30) |
| R7 | 0.61(2.4) | 0.39(24) |
| R8 | 0.64(2.8) | 0.36(15) |

**Table S3.** Tri-exponential fit parameters for residence time correlation function for all ions in the ‘core’ region of the coacervate in the DI water system. Fit parameters for all eight independent simulations are presented here.

| All ion - ‘core’ - DI water |  |  |  |
| --- | --- | --- | --- |
| Set No. | $a_1(\tau_1/\text{ns})$ | $a_2(\tau_2/\text{ns})$ | $a_3(\tau_3/\text{ns})$ |
| R1 | 0.38(3.2) | 0.35(83) | 0.27(1479) |
| R2 | 0.44(3.2) | 0.10(64) | 0.46(3187) |
| R3 | 0.43(2.7) | 0.21(60) | 0.36(4730) |
| R4 | 0.52(6.8) | 0.23(102) | 0.25(7237) |
| R5 | 0.55(3.8) | 0.12(41) | 0.33(5854) |
| R6 | 0.57(3.7) | 0.14(48) | 0.29(3707) |
| R7 | 0.38(4.1) | 0.29(18) | 0.32(3443) |
| R8 | 0.49(2.4) | 0.36(33) | 0.15(260) |

**Table S4.** Bi-exponential fit parameters for residence time correlation function for all ions in the ‘surface’ region of the coacervate in the DI water system. Fit parameters for all eight independent simulations are presented here.

| All ion - ‘surface’ - DI water |  |  |
| --- | --- | --- |
| Set No. | $a_1(\tau_1/\text{ns})$ | $a_2(\tau_2/\text{ns})$ |
| R1 | 0.70(5.2) | 0.31(55.6) |
| R2 | 0.67(6.3) | 0.33(74) |
| R3 | 0.74(6.2) | 0.26(43) |
| R4 | 0.65(6.6) | 0.35(64) |
| R5 | 0.51(4.2) | 0.49(70) |
| R6 | 0.56(6.4) | 0.44(113) |
| R7 | 0.43(5.5) | 0.57(123) |
| R8 | 0.56(5.4) | 0.44(81) |

**Table S5.** Tri-exponential fit parameters for residence time correlation function for *Na*^+^ ions in the ‘core’ region of the coacervate in the supernatant system. Fit parameters for all eight independent simulations are presented here.

| $Na^+$ ion - ‘core’ - supernatant | | | |
| --- | --- | --- | --- |
| Set No. | $a_1(\tau_1/\text{ns})$ | $a_2(\tau_2/\text{ns})$ | $a_3(\tau_3/\text{ns})$ |
| R1 | 0.23 (3.6) | 0.62 (35) | 0.16 (724) |
| R2 | 0.44 (4.5) | 0.23 (166) | 0.33 (9372) |
| R3 | 0.42 (4.6) | 0.36 (41) | 0.22 (2305) |
| R4 | 0.38 (5.3) | 0.38 (47) | 0.25 (7597) |
| R5 | 0.43 (4.8) | 0.17 (68) | 0.40 (7896) |
| R6 | 0.41 (7.1) | 0.45 (61) | 0.14 (3252) |
| R7 | 0.47 (4.0) | 0.53 (35) | — |
| R8 | 0.30 (2.4) | 0.54 (23) | 0.17 (245) |

**Table S6.** Bi-exponential fit parameters for residence time correlation function for *Na*^+^ ions in the ‘surface’ region of the coacervate in the supernatant system. Fit parameters for all eight independent simulations are presented here.

| $Na^+$ ion - ‘surface’ - supernatant | | |
| --- | --- | --- |
| Set No. | $a_1(\tau_1/\text{ns})$ | $a_2(\tau_2/\text{ns})$ |
| R1 | 0.44 (2.3) | 0.31 (55.6) |
| R2 | 0.43 (3.6) | 0.33 (74) |
| R3 | 0.33 (1.8) | 0.26 (43) |
| R4 | 0.45 (3.0) | 0.35 (64) |
| R5 | 0.36 (2.7) | 0.49 (70) |
| R6 | 0.40 (2.9) | 0.44 (113) |
| R7 | 0.47 (4.0) | 0.53 (35) |
| R8 | 0.42 (2.3) | 0.44 (81) |

**Table S7.** multi-exponential fit parameters for residence time correlation function for *Na*^+^ ions in the ‘core’ region of the coacervate in the DI water system. Fit parameters for all eight independent simulations are presented here.

| $Na^+$ ion - 'core' - DI water | | | |
| --- | --- | --- | --- |
| Set No. | $a_1(\tau_1/\text{ns})$ | $a_2(\tau_2/\text{ns})$ | $a_3(\tau_3/\text{ns})$ |
| R1 | 0.22 (7.6) | 0.44 (92) | 0.35 (1564) |
| R2 | 0.25 (4.7) | 0.11 (108) | 0.64 (3430) |
| R3 | 0.24 (5.2) | 0.26 (71) | 0.49 (5073) |
| R4 | 0.46 (14) | 0.30 (159) | 0.24 (32210) |
| R5 | 0.20 (2.7) | 0.15 (59) | 0.65 (6139) |
| R6 | 0.12 (3.3) | 0.25 (55) | 0.63 (3768) |
| R7 | 0.19 (4.3) | 0.13 (40) | 0.68 (2119) |
| R8 | 0.37 (5.9) | 0.63 (64) | — |

**Table S8.** Bi-exponential fit parameters for residence time correlation function for *Na*^+^ ions in the ‘surface’ region of the coacervate in the DI water system. Fit parameters for all eight independent simulations are presented here.

| $Na^+$ ion - ‘surface’ - DI water | | |
| --- | --- | --- |
| Set No. | $a_1(\tau_1/\text{ns})$ | $a_2(\tau_2/\text{ns})$ |
| R1 | 0.40 (6.1) | 0.60 (62) |
| R2 | 0.44 (7.8) | 0.56 (87) |
| R3 | 0.48 (7.2) | 0.52 (49) |
| R4 | 0.28 (4.8) | 0.72 (70) |
| R5 | 0.21 (7.3) | 0.79 (72) |
| R6 | 0.28 (8.6) | 0.72 (117) |
| R7 | 0.43 (5.5) | 0.77 (129) |
| R8 | 0.56 (5.4) | 0.73 (83) |

**Table S9.** Single-exponential fit parameters for residence time correlation function for *Cl*^−^ ions in the ‘core’ region of the coacervate in the supernatant system. Fit parameters for all eight independent simulations are presented here.

| $Cl^-$ ion - ‘core’ - supernatant | | |
| --- | --- | --- |
| Set No. | $a_1(\tau_1/ns)$ | $a_2(\tau_2/ns)$ |
| R1 | 2.8 | – |
| R2 | 2.8 | – |
| R3 | 2.4 | – |
| R4 | 4.0 | – |
| R5 | 4.4 | – |
| R6 | 3.4 | – |
| R7 | 0.91(3.8) | 0.09 (34271) |
| R8 | 2.76 | – |

**Table S10.** Single-exponential fit parameters for residence time correlation function for *Cl*^−^ ions in the ‘surface’ region of the coacervate in the supernatant system. Fit parameters for all eight independent simulations are presented here.

| $Cl^-$ ion - 'surface' - supernatant | |
| --- | --- |
| Set No. | $a_1(\tau_1/ns)$ |
| R1 | 3.9 |
| R2 | 5.4 |
| R3 | 4.2 |
| R4 | 4.0 |
| R5 | 3.2 |
| R6 | 3.2 |
| R7 | 2.7 |
| R8 | 3.9 |

**Table S11.**
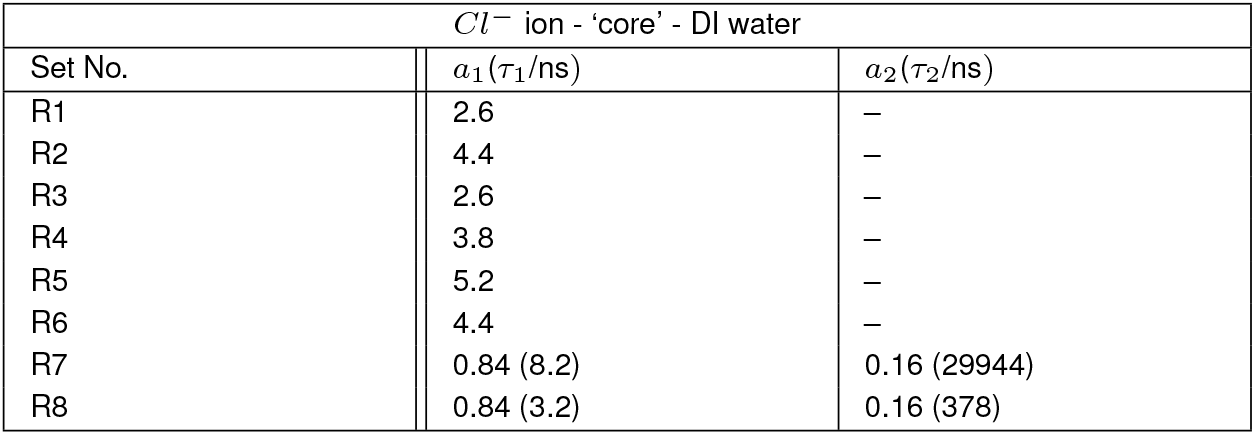
Single-exponential fit parameters for residence time correlation function for *Cl*^−^ ions in the ‘core’ region of the coacervate in the supernatant system. Fit parameters for all eight independent simulations are presented here.

| $Cl^-$ ion - 'core' - DI water | | |
| --- | --- | --- |
| Set No. | $a_1(\tau_1/ns)$ | $a_2(\tau_2/ns)$ |
| R1 | 2.6 | — |
| R2 | 4.4 | — |
| R3 | 2.6 | — |
| R4 | 3.8 | — |
| R5 | 5.2 | — |
| R6 | 4.4 | — |
| R7 | 0.84 (8.2) | 0.16 (29944) |
| R8 | 0.84 (3.2) | 0.16 (378) |

**Table S12.** Single-exponential fit parameters for residence time correlation function for *Cl*^−^ ions in the ‘surface’ region of the coacervate in the supernatant system. Fit parameters for all eight independent simulations are presented here.

| $Cl^-$ ion - ‘surface’ - DI water | |
| --- | --- |
| Set No. | $a_1(\tau_1/ns)$ |
| R1 | 6.5 |
| R2 | 8.9 |
| R3 | 6.8 |
| R4 | 8.5 |
| R5 | 4.4 |
| R6 | 6.7 |
| R7 | 4.6 |
| R8 | 5.1 |

## Supporting Movies

**Movie S1.**
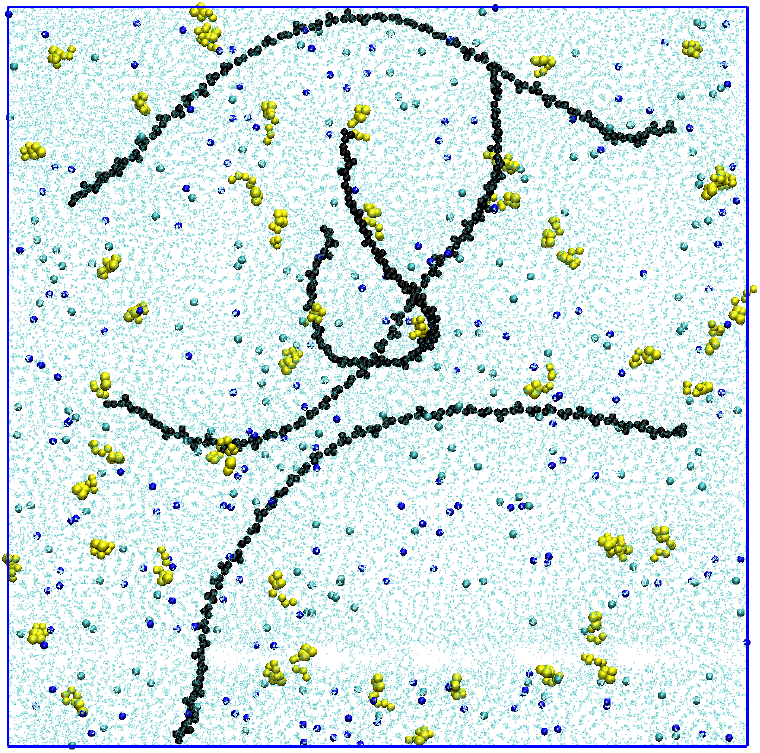
**Movie of the coacervation process starting from a completely dispersed phase of an aqueous mixture of 4 PDDA chains of 50 monomers each and 50 ATP molecules in the supernatant (i.e**., **in the presence of 200 Na^+^ and 200 Cl**^−^ **counterions). Black and yellow beads represent PDDA chains and ATP molecules, respectively. Blue and cyan beads are Na**^+^ **and Cl**^−^ **ions, respectively. Water molecules are shown as cyan dots for visual clarity. Each frame is labeled with the simulation time for the corresponding snapshot. At the simulation time of 3.5** *μ***S PDDA-ATP coacervate along with associated counterions and water molecules (purple colored points shown only for that frame) are transferred to DI water. Then simulation in DI water is shown with frames labelled with corresponding simulation times**.

